# Genome-wide screening in human kidney organoids identifies novel aspects of nephrogenesis

**DOI:** 10.1101/2021.05.26.445745

**Authors:** Rosemarie Ungricht, Laure Guibbal, Marie-Christine Lasbennes, Vanessa Orsini, Martin Beibel, Annick Waldt, Rachel Cuttat, Walter Carbone, Anne Basler, Guglielmo Roma, Florian Nigsch, Jan Tchorz, Dominic Hoepfner, Philipp S. Hoppe

## Abstract

Human organoids allow studying proliferation, lineage specification, and three-dimensional tissue development. Due to the inherent multicellular complexity, interrogation by systematic genetic methodologies is challenging. Here, we present the first genome-wide CRISPR screen in iPSC-derived kidney organoids. The combination of genome editing, longitudinal sampling and sorting of specific cell populations enabled us to uncover novel biology from development to tubular morphogenesis. We validated the high quality of our screen by individual hit follow up and comparisons to kidney disease datasets. The discovery of a novel regulatory mechanism which controls epithelial proliferation via a trans-activating but cis-inhibitory effect of the Notch ligand Jagged1 proves how mosaic knockouts generated by pooled CRISPR screening in organoids can identify novel ways of communication between heterogeneous cell populations in complex tissues. Collectively, these data demonstrate the feasibility of using complex iPSC- derived organoids for genome-scale screening and serves as a benchmark for future organoid CRISPR screens.

## Introduction

In recent years, system-scale methodologies have enabled the identification of intricate cellular signaling networks and a pleiotropy of cellular states in model systems (Doench, 2018). However, knowledge of how cellular behavior is controlled in the development of human tissues and in response to disturbances in their spatio-temporal context is limited. Organoid models recapitulate three-dimensional tissue structures *in vitro* and thus provide an opportunity to deploy large-scale functional genetics beyond their application in cancer (Michels et al., 2020; Ringel et al., 2020) to dissect lineage commitment and morphogenesis.

Starting from human pluripotent stem cells (iPSCs), directed differentiation protocols can recapitulate embryonic development and result in complex, self-organizing structures of numerous tissues (Dutta et al., 2017). In recent years, several stepwise protocols have been published which recapitulate kidney development by the generation of renal progenitor cells in a dish. These progenitor cells ultimately self-organize to form hundreds of nephron-like structures containing podocytes, epithelial tubules and interstitial cell types (Freedman et al., 2015; Morizane and Bonventre, 2017; Przepiorski et al., 2018; Taguchi and Nishinakamura, 2017; Takasato et al., 2016). Extensive characterization of marker expression and structural patterning by microscopy revealed the elaborate tissue morphology it is possible to generate within these kidney organoids, including encapsulated glomeruli and properly segmented and oriented epithelial tubules containing ultrastructural features such as a brush border (Morizane et al., 2015; Takasato et al., 2015). Organoid composition between methodologies has been analyzed and compared in multiple studies by single cell RNA sequencing (scRNA-seq). These studies all conclude organoid models to be a close resemblance to human fetal kidney whilst also providing large datasets describing the different cell populations generated by the various protocols for further interrogation (Combes et al., 2019; Czerniecki et al., 2018; Phipson et al., 2019; Wu et al., 2018).

Inherited kidney diseases comprise at least 10% of adult and nearly all cases of renal-replacement therapy in children. Currently more than 160 rare kidney diseases have been mapped genetically (Devuyst et al., 2014). A number that is increasing steadily with whole exome sequencing (WES). Here, stem-cell derived kidney organoids raise the exciting prospect of offering a scalable, human cellular system suited to model these genetic diseases, either using patient-derived iPSCs or CRISPR/Cas9 genome editing to induce the specific mutations. Human organoids which model disease can complement *in vivo* experiments, increase throughput to allow comparison of pathophysiological mechanisms whilst also giving the opportunity of personalized drug screening. A number of monogenetic kidney diseases have already been successfully modelled in kidney organoids (Cruz et al., 2017; Forbes et al., 2018; Hale et al., 2018; Lindgren et al., 2017; Low et al., 2019; Przepiorski et al., 2020). As these models continue to advance in complexity, their potential will also increase and would profit from a systems-scale view on patho-phenotypes that can be modelled in these systems.

In this study we used iPSC-derived kidney organoids to assess pathways relevant in human kidney development in both health and disease by pooled genome-wide CRISPR screening. We generated a complex, functional dataset including temporal and cell-type specific resolution which allowed us to (i) benchmark the general feasibility of genome-wide screening in kidney organoids, (ii) assess the potential of human kidney organoids to model genetic disorders on a broad scale and (iii) uncover novel biology which was verified in follow-up experiments. Our screen identifies ROCK inhibition as a beneficial event in the early stages of directed differentiation towards nephron progenitor cells, a protocol modification that increases the robustness of organoid production from different stem cell lines. Moreover, our hit list spans a wide range of disease-relevant genes, particularly those associated with congenital anomalies and ciliopathies. We show that general ciliary ablation by *KIF3A* knockout (KO) or ciliary dysfunction upon heterozygous *OFD1* mutation mimics a ciliopathic renal phenotype in kidney organoids. Lastly, we describe a novel cis-inhibitory mechanism of the Notch ligand Jagged1 that controls tubular proliferation – a control mechanism needed to avoid cystic growth, in human organoids and a mouse model. Taken together, we provide herein a resource of genes that are functionally important in human kidney development in both health and disease, paving the way for future disease-specific whole-genome screens in kidney or other organoid models.

## Results

### Setting up the protocol for pooled CRISPR screening in kidney organoids

To determine the optimal conditions for a genome-wide CRISPR screen, we initially characterized i) organoid growth rate heterogeneity to identify potential biases, ii) organoid composition to establish a sorting regime and iii) organoid development to define key time points. Taking the advantage of being fully chemically defined for eventual upscaling efforts, we used the protocol described by Morizane and Bonventre (2017) to efficiently generate nephron-like structures from human iPSCs within 3 weeks (Figure 1A). Unlike other organoid systems that grow from single adult stem cells (e.g. intestinal organoids), kidney organoids are established by self-aggregation of high numbers (1000s) of nephron progenitor cells (NPCs) and thus may avoid the issue of clonal growth. Time-lapse imaging demonstrated highly synchronized growth of individual organoids (Figure 1B). Unsupervised clustering of 24’055 single cell transcriptomes from days 21, 28 and 35 revealed a total of 11 clusters that could be attributed to epithelial, podocyte or stromal lineages (Figure 1C). Based on previously published scRNA-seq datasets (Wu et al., 2018; Young et al., 2018) the clusters identified indicated appropriate kidney-specific differentiation (Table S1). Furthermore, we confirmed EPCAM as a suitable marker to sort all epithelial subtypes (proximal tubule, loop of henle and distal tubule) for screening (Figure 1D). The high degree of consistency between replicates and published scRNA-seq data using the same protocol suggested stability even between different laboratories (Figure S1A, B) (Wu et al., 2018). Between day 21 and 35, we observed an upregulation of known fibrosis-associated genes (*COL1A1*, *DCN*, *ACTA2*, *LGALS1*, *TGFB1*, *MFAP4*) in stromal cells, as well as an upregulation in epithelial cell genes (*IGFBP7*, *JUN*, *JUNB*, *VIM*, *SOX4*, *CTGF*) suggesting a fibrotic response and initiation of epithelial to mesenchymal transition, confirmed by increased immunolabelling of collagen fibers (Figure S1C). Besides that, cell type identity appeared to be stable between day 28 and 35 with no further maturation of tubular epithelial cells (TECs) or podocytes nor large changes in cell type frequency distribution (Figure 1C). Similarly, flow cytometry analysis of EPCAM+ cells and cycling cells revealed a plateau after day 28 with ∼5% of TECs and ∼20% of stromal cells still cycling (Figure 1F). The first cell type specific markers became visible concurrently with the decrease in proliferation rate (Figure 1E), suggesting lineage specification occurring prior to day 14. We defined this time point to discriminate between functional contribution of genes to kidney development and proliferation within the lineage.

**Figure 1.**
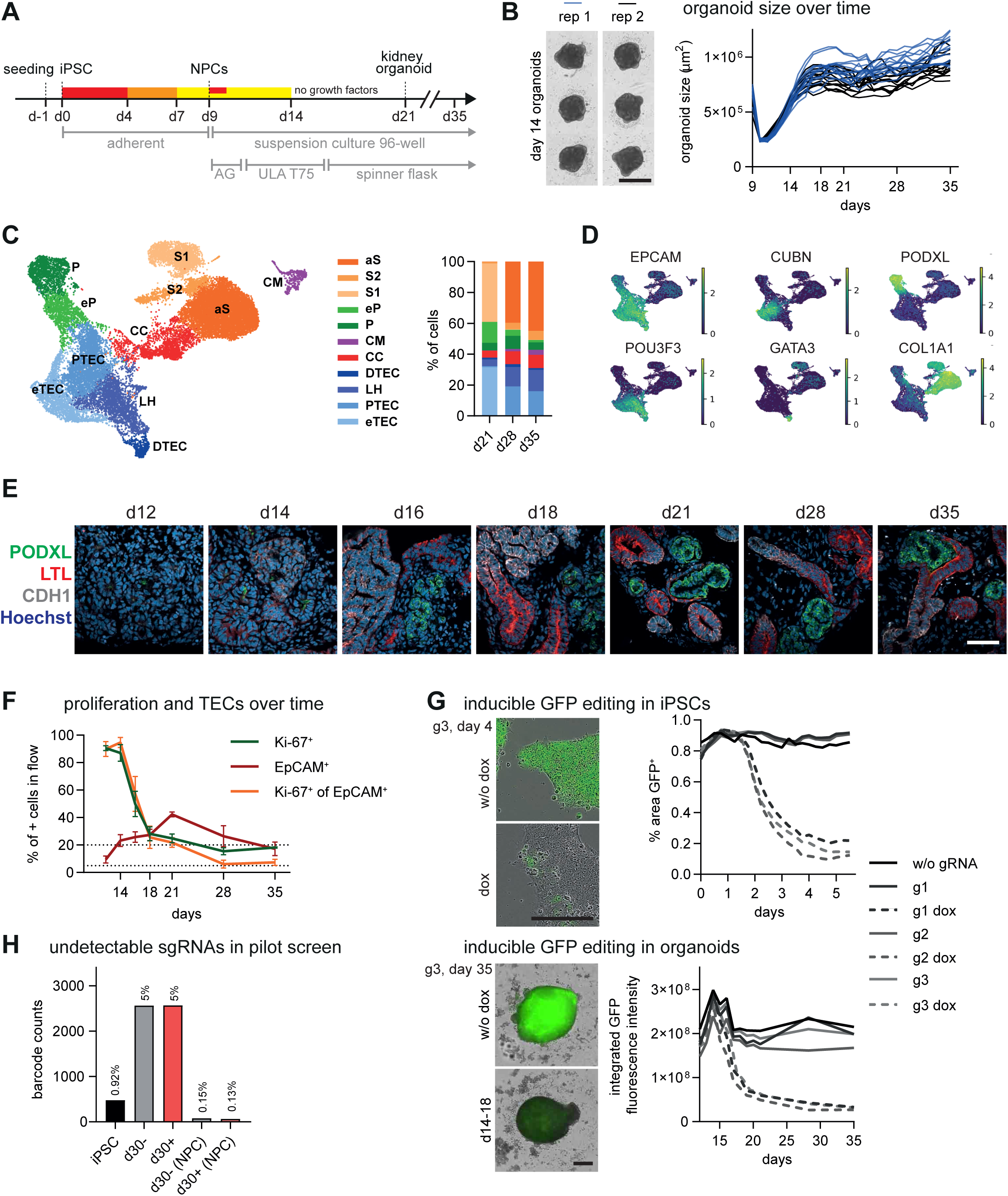
Characterization of kidney organoid system. (A) Schematic representation of differentiation protocol for kidney organoid production in small and large scale. NPCs: nephron progenitor cells; AG: Aggrewell. (B) Example brightfield images of organoids at day 14. Time-dependent size tracking (beginning with formation of 3D suspension culture at day 9) demonstrates consistent growth of individual organoids. N=2; n=14. Scale bar 1 mm. (C) Single cell RNA-seq. UMAP showing cell populations identified by unsupervised clustering of cells with similar transcriptomes for all time points (d21, d28 and d35). Stacked bar plot with frequency distribution of cell types at indicated time points. eTEC: early tubule epithelial cell; PTEC: proximal TEC; LH: loop of henle; DTEC: distal TEC; CC: cycling cell; CM: cap mesenchyme; P: podocyte; eP: early podocyte; S1: stroma 1; S2:stroma 2; aS: activated stroma. n=2. (D) Feature plot highlighting expression of EPCAM in tubular cells (CUBN, POU3F3, GATA3) but not in podocytes (PODXL) and stroma (COL1A1). (E) Time-course of immunostained cryosections of kidney organoids display expression of podocyte (PODXL) and tubular markers (LTL and CDH1) starting between day 14 and 16. Concomitantly, nephrons begin to pattern. Representative images of N=3 individual experiments. Scale bar 50 µm. (F) Flow cytometry single cell-analysis of organoids dissociated at indicated time points to quantify cycling cells (Ki-67^+^) and cycling tubular cells (Ki-67^+^ of EPCAM^+^). Dashed lines indicate 5 and 20 percentile. N=3, 8 organoids each. (G) Evaluation of inducible genome editing in iPSCs and organoids. eGFP expressing iPSCs, transduced and selected for the integration of lentivirus harboring sgRNAs against eGFP were either kept in culture without induction of Cas9 expression (solid lines) or induced by dox (dashed lines). Editing was induced in iPSCs or organoids from day 14 to 18. Scale bar 300 µm. (H) Pilot screen sgRNA representation analysis. Organoids derived from cells infected with the lenti-CRISPR library at the iPSC (sample iPSC, infection + 3 passages) or NPC stage, both at coverage 50 cells/gRNA, were edited (+dox d14-18; +) or unedited (-) and collected at day 30 (d30) to assess gRNA representation by NGS. Numbers above bars indicate percentage of missed sgRNA compared to total library.

To study key processes in kidney (organoid) development we combined longitudinal sampling and timed, inducible genome editing (see below, Figure 2A). We generated an inducible Cas9 iPSC line (iCas9) harboring a single copy of a doxycycline (dox) inducible system including an insulator to prevent leaky expression, integrated into the *AAVS1* locus (Ihry et al., 2018). In the absence of dox, three cell lines transduced with a lentivirus delivering constitutively expressed sgRNAs targeting essential genes (*RAN*, *PLK1* and *POLR2B*, respectively) grew the same as wildtype (WT) control, showing tight control over Cas9 expression. Induction of Cas9 (+dox) elicited an immediate growth arrest and led to massive cell death (Figure S2A). Editing efficiency was also assessed using an eGFP expressing line to rule out that the observed growth arrest stems from the induction of double strand breaks *per se* which induces a certain degree of toxicity in iPSCs (Haapaniemi et al., 2018; Ihry et al., 2018). In agreement with our first results, sgRNAs targeting eGFP led to an efficient reduction of eGFP within 4 days of dox addition in iPSCs and organoids (Figure 1G, Figure S2B, C).

**Figure 2.**
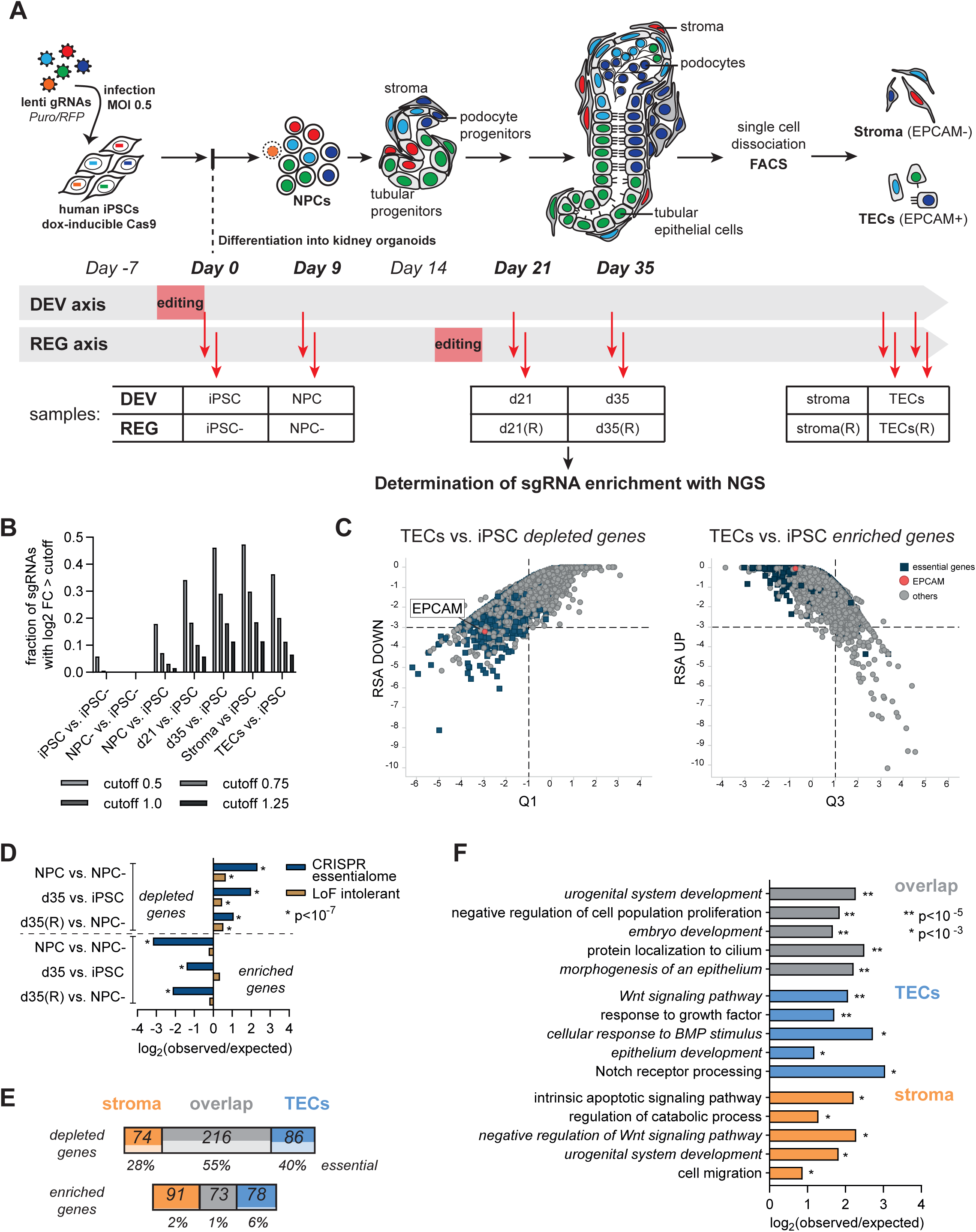
Pooled CRISPR screening in kidney organoids during development. (A) Experimental outline of pooled genome-wide CRISPR screen(s) in kidney organoids. A genome-scale lentiviral sgRNA library was transduced in iPSCs at an MOI of 0.5 followed by selection with puromycin. Cultures we split for two editing regimes during differentiation: Editing induced in iPSCs (DEVelopment axis) or after day 14 (REGeneration axis). Pictures illustrate that single organoids contain multiple gRNAs. End-stage organoids (day 35) were dissociated and tubular cells or stromal fractions sorted by FACS. Stages at which samples were collected are outlined. (B) Fraction of sgRNA with absolute log2 fold change > cutoff over time. Note that no sgRNAs score in unedited NPCs suggesting the absence of a bottleneck in early differentiation. See also Figure S3C. (C) RSA plot for TECs compared to iPSCs in DEV axis of the screen. Blue squares represent essential genes identified by CRISPR screening in cancer cell lines (Hart et al., 2015). Lines indicate cutoff for stringent hit list (FDR < 1%, Table S3). Note the depletion of EPCAM, the epitope sorted for to enrich tubular cells (positive control). (D) Enrichment plot for essential genes, CRISPR essentialome as in (C) or LoF intolerant genes (pLI 0.9-1) from the ExAC dataset (Lek et al., 2016) in depleted or enriched genes from DEV and REG axis (cutoff RSA <-2). (E) Hit distribution in TECs, stroma and overlap, all compared to iPSC with stringent cutoff of RSA <- 3. Percentages indicate essential genes (Hart et al., 2015). (F) Enriched GO terms among genes scoring in TECs only, stroma only and overlap from (E). Essential genes were excluded. Source: ToppGene suite.

Next, we conducted a pilot screen to identify potential bottlenecks during the differentiation from iPSCs to NPCs to kidney organoids. With a low coverage of 50 cells per barcode and a MOI of 0.5, a library of 51’644 gRNAs was transduced at iPSC or NPC stage, followed by differentiation in the presence or absence of genome editing. This pilot revealed a bottleneck in the first 9 days of differentiation, as 5% of the library could not be recovered even in the absence of genome editing (Figure 1H). Transduction of NPCs at 50 cells per gRNA did not lead to any loss of barcodes, corroborating the stability of differentiation from the NPC stage onwards. For the genome-wide screen we incorporated these learnings and increased coverage to at least 300 cells per gRNA at all stages after transduction of iPSCs and included a ‘bottleneck’ control (without dox) up to the NPC stage.

### Genome-scale CRISPR screening in organoids for development and regeneration

To enable an efficient loss-of-function screening process, we leveraged a genome-wide lentiviral CRISPR library (18360 genes, 5 sgRNAs per gene) investigating two paradigms: The first to study functional contribution to kidney development, the second to capture genes involved in cell proliferation within a lineage. For the first experimental arm, editing was induced in iPSCs and gRNA representation assessed over time in bulk (iPSCs, NPCs, day 21 and day 35 whole organoids), tubular (EPCAM^+^) or non-epithelial (stroma, EPCAM^-^) cell fractions following dissociation and cell sorting by flow cytometry (Figure 2A, DEVelopmental axis). The EPCAM^-^fraction predominantly contains stromal cells as podocytes fall into an EPCAM weak gate which was excluded (thus called ‘stroma’ from now on, Figure S3A). In the second arm, editing was induced after the last lineage choice at day 14 using the same sampling and sorting regime (REGeneration axis). This arm served in addition as a technical control for sampling bottlenecks in the critical early differentiation phase.

Changes in sgRNA representation identified by barcode sequencing are related to the number of cell duplications. Based on cell counting, we estimate an average of 5 population doublings from iPSCs to NPCs. Despite this large increase in cell number, in the absence of editing no sgRNAs scored significantly (Figure 2B), very few barcodes were lost (0.15% not detected) in NPCs (Figure S3B) and the fold change distribution between NPC and iPSC was flat (Figure S3C), demonstrating the general feasibility of our approach. At day 9 NPCs were aggregated into 14400 organoids (20000 cells per organoid) per axis and grown in spinner flasks from day 16 onwards, carefully monitored over time for marker expression and 3D patterning (Figure S3D&E). In agreement with more cell duplications, identity changes, and the longer culturing time, more sgRNAs scored over time in the DEV axis (Figure 2B, S3B). To identify relevant processes on the gene level, sgRNA numbers were processed by the redundant small interfering RNA activity (RSA) test and plotted against first or third quartile Z-score (König et al., 2007) (see also methods and gene ranks in Table S2).

As expected, EPCAM (being used to sort TECs) scored among the top depleted genes in TECs in both axes (for example rank 169 in TECs vs iPSC, DEV axis; *p* = 0.005), suggesting successful execution of the screen incl. KO, sorting and processing of the sgRNA counts (Figure 2C). Next, we benchmarked the performance of our screen by comparing hits to core-essential genes defined by CRISPR screening (CRISPR essentialome, (Hart et al., 2015)) and loss-of-function (LoF) intolerant genes (Lek et al., 2016). KO of genes essential for survival of cells *per se*, should lead to depletion of the corresponding gRNAs during CRISPR screening. Indeed, already at the NPC stage essential genes significantly dropped out. Essential genes were significantly enriched in dropout hits (for example 453 genes in 988 top hits in RSA DOWN NPC (edited) vs. NPC^-^ (unedited) are ‘CRISPR essentialome genes’, *p* = 1.25×10^-244^) and almost not present in overrepresented genes (9/900 in RSA UP NPC vs. NPC^-^, Figure 2D *enriched genes*). Essential and LoF intolerant genes remained a large fraction of the depleted genes until day 35, in sorted cell types and in the REG axis (Figure 2D and blue squares in Figure 2C for TECs, RSA DOWN). The strong representation of essential genes in our screen provided another control parameter and corroborated our approach.

### Pathways contributing to kidney development

To functionally dissect and infer pathway dependence, we ranked hits for genes specifically enriched or depleted in TECs and/or stromal cells. We used stringent cutoff criteria for this first pass analysis with a false discovery rate (FDR) of less than 1% (Figure 2E, Table S3; RSA <-3; further referred to as Hit list 1). As expected, a large proportion again fell into the core-essential genes category – with up to 55% essential genes in hits depleted both in TECs and stroma. These were excluded from further analysis. The residual 435 hits were grouped by cell type and assessed for enrichment of biological pathways and processes (Chen et al., 2009). Overlapping TEC-specific and stromal hits were all significantly associated with kidney development-related GO terms among the top associated biological processes, including ‘urogenital system development’, ‘embryonic development’ and ‘epithelium development’ (Figure 2F). In good agreement with the inductive chemical cues used to differentiate human stem cells into kidney organoids, BMP and Wnt signaling-related terms were also enriched. In addition, our data confirmed a link between tubular specification and Notch signaling (with *RBPJ, PSENEN and ADAM10* depleted in TECs) and substantiated the importance of Sonic Hedgehog signaling for early kidney development (with *PTCH1, SUFU, GLI3* plus 21 genes connected to primary cilia strongly enriched in both cell types) (see also Figures 5, 7). In total, 20% of the queried top hits were associated to known kidney-related biological processes (GO terms in *italic* Figure 2F). Another expected hit family are tumor suppressor genes, which were overrepresented in both TECs and stroma and associated with the term ‘negative regulation of cell population proliferation’ (22/27 enriched in RSA UP; among them e.g. *TSC1*, *TSC2*, *WT1*, *NF1*, *RB1* and *PTEN)*.

### Genome-wide screening can improve early steps in specification towards the kidney lineage

Inducible editing and longitudinal sampling can provide insight on the contribution of individual cellular pathways during the differentiation process. Hence, hits can be subdivided into three families: (1) genes required for early metanephric mesenchyme induction (Figure 3A, green), (2) genes involved in nephrogenesis (orange) and (3) genes influencing maturation (yellow). We generated hit lists for these three categories using a cutoff of RSA<-2, which corresponds to a FDR of less than 5%. Hits are further attributed according to cell type (TECs, stroma or overlap) and effect direction (enriched or depleted) (Table S4; Hit list 2).

**Figure 3.**
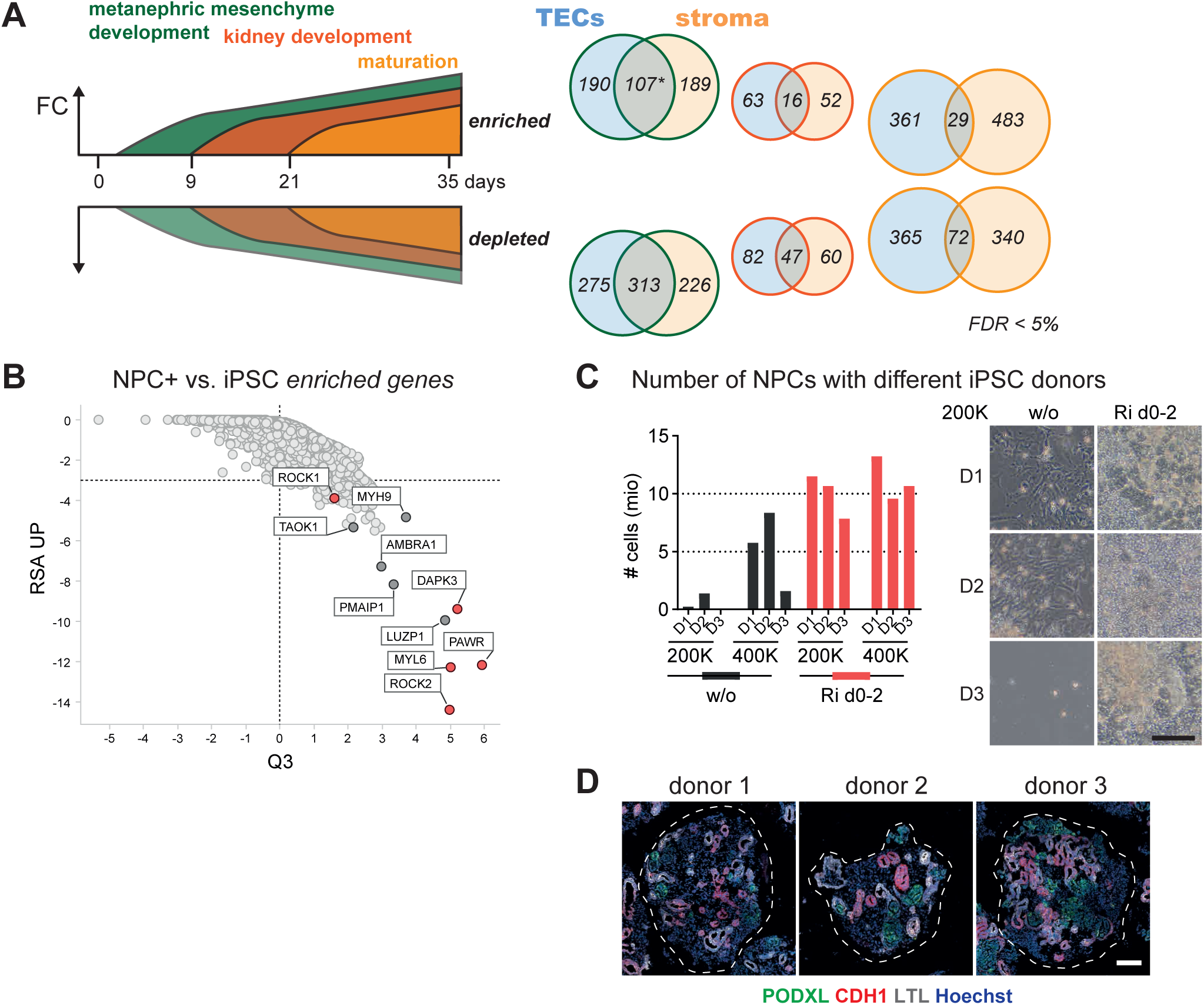
Improving early steps towards kidney lineage specification using ROCKi. (A) Scheme illustrating how time point sampling allows discrimination between early, intermediate and late functional contribution of genes. Analysis over time together with endpoint sorting of TECs and stroma allows to bucket hits (RSA <-2 and FDR <5%, w/o essential genes) into 18 categories. Numbers in Venn diagrams indicate number of hits in buckets. *: category in which genes linked to the Rho/ROCK pathway fall. (B) Gene KOs that get enriched in NPCs (DEV axis). Marked in red is the STRING network around ROCK1/2. (C) Treatment of cells with ROCKi during the first 2 days of differentiation leads to pronounced increased survival, as assessed by absolute cell number at day 9 of differentiation. Effect is more apparent at lower initial seeding density (200K: 200000 cells seeded per 6-well). Donor 1 (D1) is our standard iPSC line. Difficult to differentiate iPSC lines (D2 and D3) react to ROCK inhibition and survive until the NPC stage. Representative brightfield images illustrate successful induction and initial 3D structuring of NPCs at day 9. Scale bar 100 µm. (D) Immunofluorescence staining on cryosections of day 21 organoids grown with ROCKi present in the first 2 days of differentiation, 200000 cells seeded initially. Scale bar 100 µm.

In the genes required for metanephric mesenchyme induction (a bucket of 107 genes), a cluster linked to Rho-associated kinases *ROCK1* and *ROCK2* (including *DAPK3*, *MYL6* and *PAWR*) scored significantly (*p* < 0.002 for all; Figure 3B). ROCK inhibition by the small molecule compound Y27632 is regularly used to increase survival of iPSC cultures when split as single cells, but can be omitted the day after seeding without influence on survival or growth rate. In our protocol, we also use ROCK inhibition when seeding iPSCs the day before differentiation (day -1 to 0). Beyond that, treatment with the ROCK inhibitor (ROCKi) Y27632 during the first 2 days of differentiation strongly increased cell survival (Figure S4). This was even more pronounced at lower seeding densities, at which cells under normal conditions do not survive the inductive cues. Longer or later treatments did not further influence differentiation. To test whether ROCKi may alleviate inter-organoid variability between different iPSC donors (for kidney see Phipson et al. (2019)), two additional iPSC lines that were challenging to bring through the first phase of differentiation were cultured with and without ROCKi during mesoderm induction. The presence of ROCKi during the first 2 days made a strong difference in the number of NPCs generated at day 9 (Figure 3C). We managed to produce well-structured organoids for all 3 iPSC lines in a parallel effort without changing any inhibitor/growth factor or seeding concentrations. This protocol modification may provide a critical advantage for the investigation of kidney organoids from different donors (Figure 3D).

### Screening hits cover kidney disease genes

To continue our analysis, our stringent Hit list 1 (Figure 2E, Table 3) was further examined for human disease associations. The most highly enriched human phenotype associations were related to kidney diseases with a reported embryonic manifestation. We observed the strongest enrichment when comparing to genes that affected both tubular and stromal cells - covering overgrowth syndromes like ‘renal cysts’, ‘enlarged kidneys’ and ‘renal neoplasm’, in addition to kidney malformations such as ‘renal dysplasia’, general ‘abnormality of the kidney’, ‘nephronophthisis’ and even ‘chronic kidney disease’ (*p* < 0.005 for all terms) (Figure 4A). The same terms associated with the larger Hit list 2 generated with less stringent cutoff criteria (RSA<-2, FDR<5%, Table S4). We can thus nominate more than 490 disease-associated genes that give a phenotype in human kidney organoids (Table S5 summarizes terms and associated genes).

**Figure 4.**
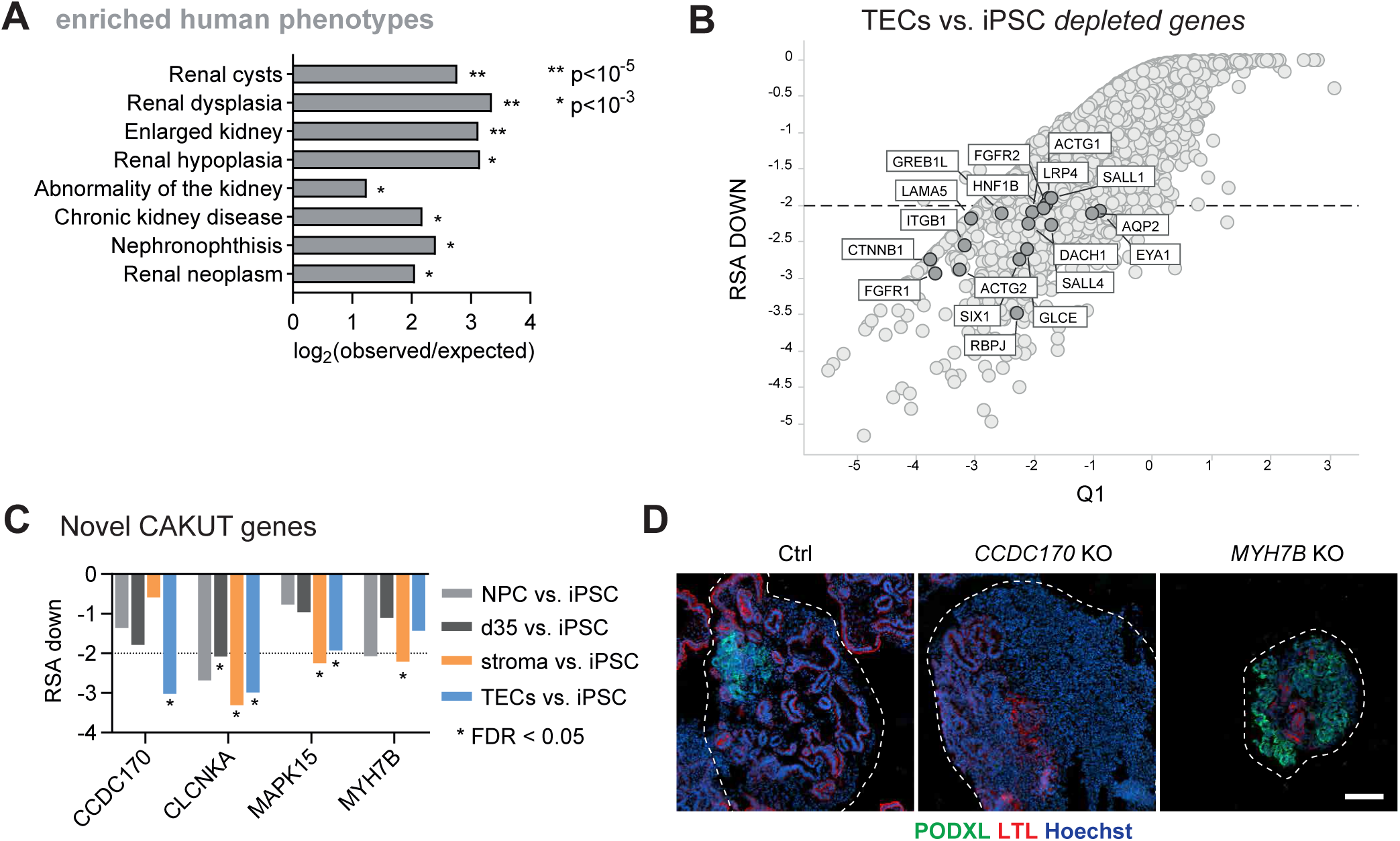
Whole-genome organoid screen identifies kidney disease-relevant genes. (A) Enriched human phenotype associations in overlapping screening hits from tubular and stromal cells (see Figure 2E). Source: ToppGene suite. A list of all genes that associate with kidney-related human phenotypes and score in the screen can be found in Table S5. (B) Genes depleted in TECs vs. iPSCs (DEV axis) that are known to cause anomalies in kidney development (human and mouse CAKUT). Essential genes were excluded in RSA plot. (C) Candidate CAKUT genes (van der Ven et al., 2018a) that score in CRISPR screen DEV axis. Note that permutation analysis (FDR) was only calculated for d35 whole organoid, stroma and TEC comparisons. (D) Immunofluorescence staining on whole organoid section of organoids at day 21 reveal less nephron tubules upon *CCDC170* KO and a lack of stroma upon *MYH7B* KO. Scale bar 100 µm.

Kidney organoids are well suited to model developmental diseases in particular due to their neonatal characteristics. The broader term ‘congenital anomalies of the kidney and urinary tract’ (CAKUT) summarizes a broad spectrum of clinical conditions with renal manifestation in early childhood that are caused by monogenic mutations leading to pathogenic dysregulation of renal morphogenesis (van der Ven et al., 2018b). CAKUT genes that affect metanephric mesenchyme morphogenesis *per se*, without crosstalk to the ureteric bud or involvement of the lower urinary tract, could score in our screen. Indeed, we identified a number of bona fide CAKUT genes including *HNF1B*, *EYA1, SALL1* and *SIX1* that were depleted in TECs, resembling a tubular agenesis phenotype (Figure 4B). When comparing our screening data to a broader list of CAKUT genes from human and mouse models (modified from Heidet et al. (2017)), a greater number of genes overlapped which gave phenotypes either in TECs or whole organoids (Figure 4B and Table S6). A study on a large cohort of 232 families with CAKUT nominated 19 single novel candidate genes that are potentially causative for monogenic CAKUT, without being able to further substantiate their relevance with additional families or experimental evidence (van der Ven et al., 2018a). 4 out of these 19 genes in our screen resulted in a relevant KO phenotype being essential for either TEC and/or stromal development: *CCDC170* dropped out late in tubular cells only (see Figure 4C for effect strength). *CLCNKA* LoF led to early depletion in NPCs and was essential for general development. *MAPK15* had a later effect on development and affected both stroma and TECs. *MYH7B* dropped out in NPCs and the stroma. Follow up on the two weaker, cell-type specific effects by individual CRISPR KO confirmed a role for CCDC170 in tubulogenesis and MYH7B in stromal growth resulting in normally sized organoids containing a disproportional amount of stroma (for *CCDC170* KO) or smaller organoids with large amounts of podocytes (*MYH7B* KO) (Figure 4D). Thus, we suggest that our *in vitro* screening data can be used to identify and prioritize novel monogenic causes of CAKUT. In addition, the organoid model can be exploited to understand the molecular mechanism of CAKUT etiology.

### Human kidney organoids model ciliopathies

Within the genes associated with kidney-related human phenotypes, we noted that a remarkable number was linked to primary cilia, either constituting ciliary components or contributing to cilia assembly. A comparison to 187 established ciliopathy genes (Reiter and Leroux, 2017) revealed a striking overrepresentation (in total 49 genes) within hits causing overproliferation of TECs and stroma (Figure 5A). 37 out of these 49 genes have been described in patients with renal phenotypes (Table S7). Consequently, our screen nominates the residual 12 genes and another 84 genes that localize or function in primary cilia as potentially causative for renal ciliopathies (Table S8).

**Figure 5.**
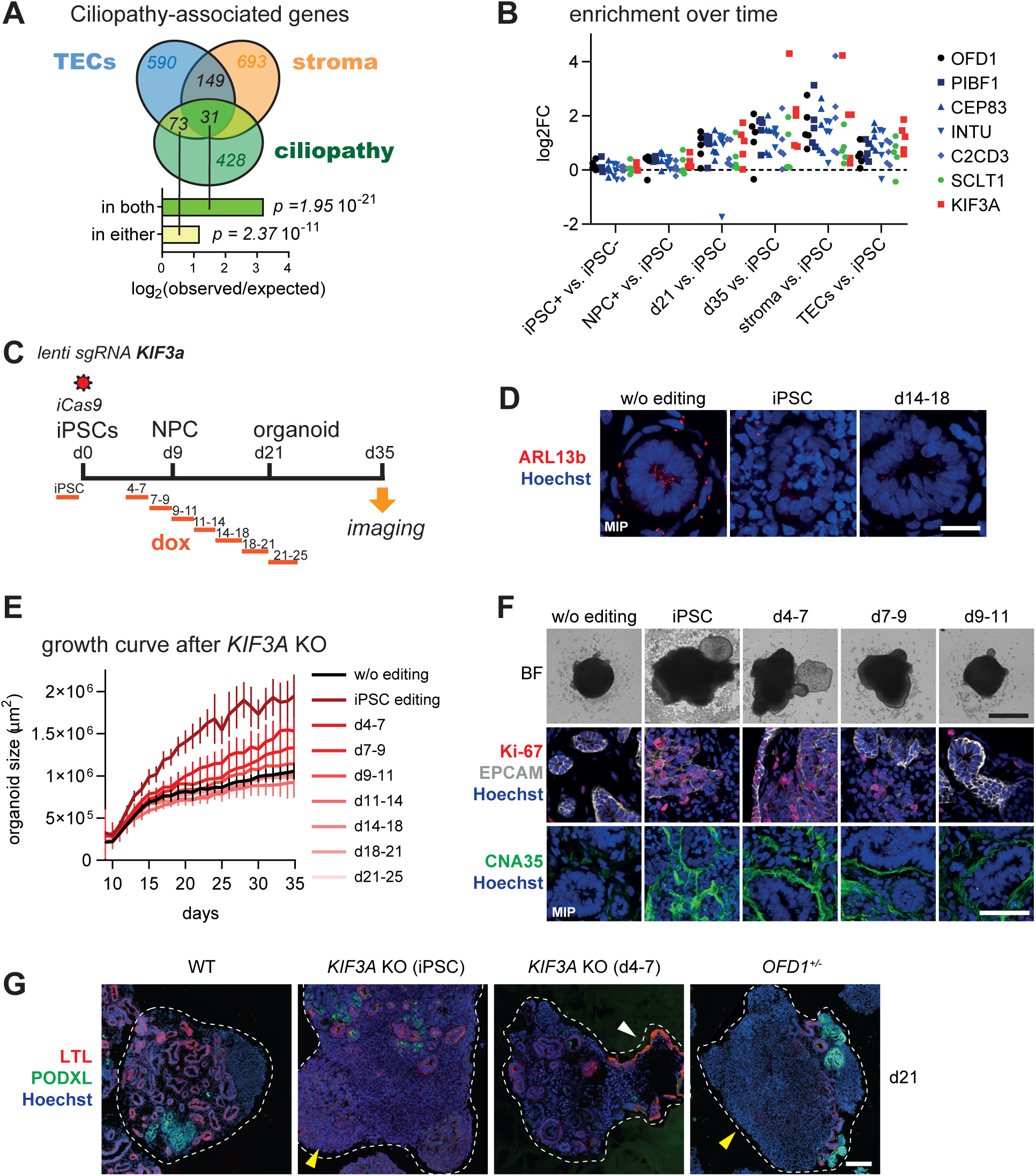
Kidney organoids allow modeling of ciliopathies in phenotype and kinetics. (A) Enrichment plot for established (187) and candidate (241) ciliopathy genes (in total 428) (Reiter and Leroux, 2017) in hits (cutoff RSA <-2; DEV axis) leading to overrepresentation of gRNAs in TECs and/or stroma. See also Table S7. (B) Example of log2FC enrichment at sgRNA level over time of 5 established ciliopathy-causing genes, *KIF3A* and *SCLT1*. (C) Experimental setup for *KIF3A* KO in iPSCs or at different stages during organoid development using inducible genome editing. (D) Immunofluorescence staining on cryosections from day 35 organoids show a complete ablation of cilia (ARL13b) after *KIF3A* KO. Scale bar 50 µm. (E) Longitudinal tracking of organoid size by brightfield imaging after *KIF3A* KO induced at variable stages during development (n=16). (F) KO in iPSC or at early stages during development leads to overgrowth and cysts as visible in time-lapse imaging. Representative brightfield images at endpoint day 35. Scale bar 1 mm. Immunofluorescence stainings on cryosections from day 35 organoids reveal misstructuring of the tubular epithelium (EPCAM), an increase in the number of cycling tubular and stromal cells (Ki-67) and collagen fiber deposition/ fibrosis (CNA35). Scale bar 50 µm. (G) Whole organoid section of polyclonal *KIF3A* KO and *OFD1^+/-^* organoids at day 21 shows stromal overgrowth (yellow arrow) and outgrowing cysts lined by a single layer of epithelial cells (LTL, white arrow). Scale bar 100 µm.

The longitudinal sampling allowed us to pinpoint in more detail when disturbance in ciliary function translated into a phenotype. Ciliary genes are enriched both in stroma and TECs from day 21 onwards, with both cell types showing comparable enrichment (Figure 5B). While the enrichment of sgRNAs over time suggested a prolonged effect on proliferation, we did observe a clear difference between both screening axis. To clarify when cilia are necessary for proper kidney development, we followed up on KIF3A, a kinesin motor reported to be essential for cilia formation and leading to cystic growth in mice if deleted in renal TECs (Lin et al., 2003). *KIF3A* KO is often used to achieve complete ciliary ablation and thus served as a good surrogate to study the role of primary cilia *per se*. We used timed, inducible editing of *KIF3A* in 3 to 4-day intervals during differentiation and followed organoid growth by live imaging (Figure 5C). With an average editing efficiency of 84.5% (not shown), we confirmed ciliary ablation by immunostaining (Figure 5D). Editing in iPSCs and early phases of differentiation (day 4-7 and 7-9) led to growth of fluid filled cysts and a striking increase in organoid size, resulting in organoids composed of 3- fold more cells at day 35 (Figure 5E&F). Notably, unlike editing before the NPC stage, the organoids with *KIF3A* KO before the NPC stage exhibited an increased growth rate particularly in the late stages of culturing (between day 18 and 35). The lack of cilia also led to severe structural abnormalities of the tubular epithelium within the organoid (see EPCAM in Figure 5F) and increased deposition of collagen fibers (see CNA35), mimicking a nephronophthisis phenotype *in vitro*. We can further confirm this phenotype in a more disease-relevant model of the X-linked oral-facial-digital syndrome (Adamiok-Ostrowska and Piekiełko-Witkowska, 2020). A heterozygous deletion of 5 bp in exon 16 of *OFD1* leading to a frameshift and premature stop (c.1898_1902del; p.A633GfsX23) results in the same kidney organoid phenotype of increased growth and mis-structuring of the nephron tubules – notably without ciliary ablation (Figure 5G and S5). As already suggested based on inducible *PKD1* deletion in mice (Lantinga-van Leeuwen et al., 2007), our results confirm the importance of an active cell cycle in tubular cells for the induction of a cystic phenotype upon ciliary dysfunction in a human system. Thus, kidney organoids are well suited to model renal ciliopathies and study molecular disease etiology.

### Pathways contributing to proliferation within the tubular lineage

Kidney regeneration after tubular damage is hypothesized to follow a similar mechanism as that which occurs during the late stages of development (Chang-Panesso and Humphreys, 2017). Therefore, we compared hits in the REG axis to hits that score late during development (Figure 6A). Comparison between the two axes increased robustness, as isolated analysis of the REG axis contained more noise due to the smaller number of populations doublings. Using this approach, we obtained a confined stringency list of 161 genes which specifically affected TECs and 122 genes which influence non-epithelial cells (Table S9). Metascape gene set enrichment analysis (GSEA) was applied for both hit lists (Zhou et al., 2019). The top enriched canonical pathways in TECs related to presenilin-1 (PS1)/Notch signaling and focal adhesion kinase (FAK) (Figure 6B). Contribution of cellular adhesion to tubular integrity also scored highly in the ‘regulation of cell adhesion’ GO term, with a large overlap in associated genes (*ARHGEF7, RASA1, ETS1*). The importance of cell adhesion and polarity was further emphasized by *CDC42*, *STK11* and *MARK2,* the key players of the epithelial polarity program that were identified in TECs in the REG axis exclusively. Another interesting hit class negatively regulates cAMP-mediated signaling. cAMP signaling is hypothesized to contribute to cystic growth of nephron epithelial cells in polycystic kidney disease (Bergmann et al., 2018; Magenheimer et al., 2006). Therefore, regulators of this pathway that impact tubular growth might be of particular interest for disease management. Noteworthy is also the coverage of 4 subunits of the chromatin modifying complex STAGA. All 4 genes scored robustly with both editing regimes and showed enrichment specifically in TECs. To the best of our knowledge, to date no functional connection between the STAGA complex and nephrogenesis has been reported and thus this constitutes a novel epigenetic angle.

**Figure 6.**
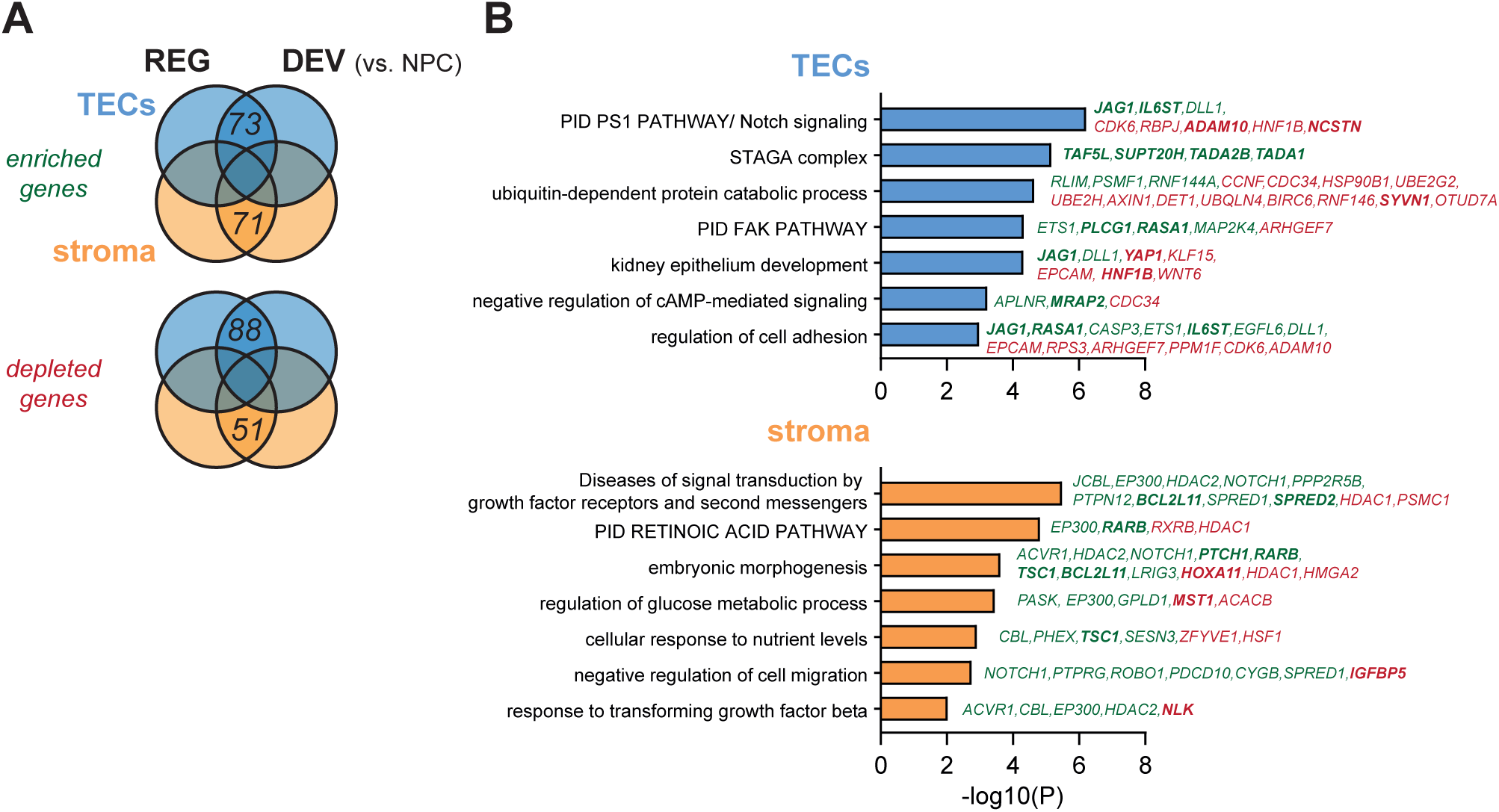
Genes influencing proliferation within kidney cell lineages. (A) Intersection of hits scoring in DEV and REG axis, specifically in TECs or Stroma. Essential genes were excluded. For cutoff see Table S9. (B) Metascape pathway enrichment for hits in (A). Genes in italic show overlap between input hit list and pathway term (green: enriched genes; red: depleted genes from screen). Selected genes that score strongly are highlighted in bold (see Table S9).

### A new role for Jag1 in restricting tubular proliferation

Besides our observation that Notch signaling is among the top enriched pathways in TECs (Figure 6B), we noticed an intriguing complex scoring pattern of Notch ligands and receptors in our data set (Figure 7A). Our screen recapitulated the necessity of Notch signaling via *NOTCH2* for tubular specification (Mukherjee et al., 2019), in that *NOTCH2*, *RBPJ*, *ADAM10*, *NCSTN* and *PSENEN* dropped out between the NPC and early organoid stage (day 21) in the DEV axis. However, these genes also scored in TECs when editing was induced after the last lineage choice, proposing a further role in tubular elongation. As our screen was conducted in a pooled setup, the finding of the Notch ligand Jagged1 as one of the strongest hits in TECs in both editing regimes is remarkable. In a genome-wide pool only few cells per organoid are edited for Jagged1, thus surrounding cells should be able to compensate for its loss. In addition, effects of ligands on neighboring cells are not read out, since affected neighbors do not contain the (Jagged1) gRNA, but any other one. Considering that Jagged1 also led to an opposing effect compared to Notch2 (more TECs upon *JAG1* KO, less TECs upon *NOTCH2* KO), the most plausible hypothesis is that Jagged1 acts in ‘cis’ within the signal sending (tubular) cell. In support of this, our scRNA-seq data revealed *JAG1* expression specifically in tubular cells (Figure S6A).

**Figure 7.**
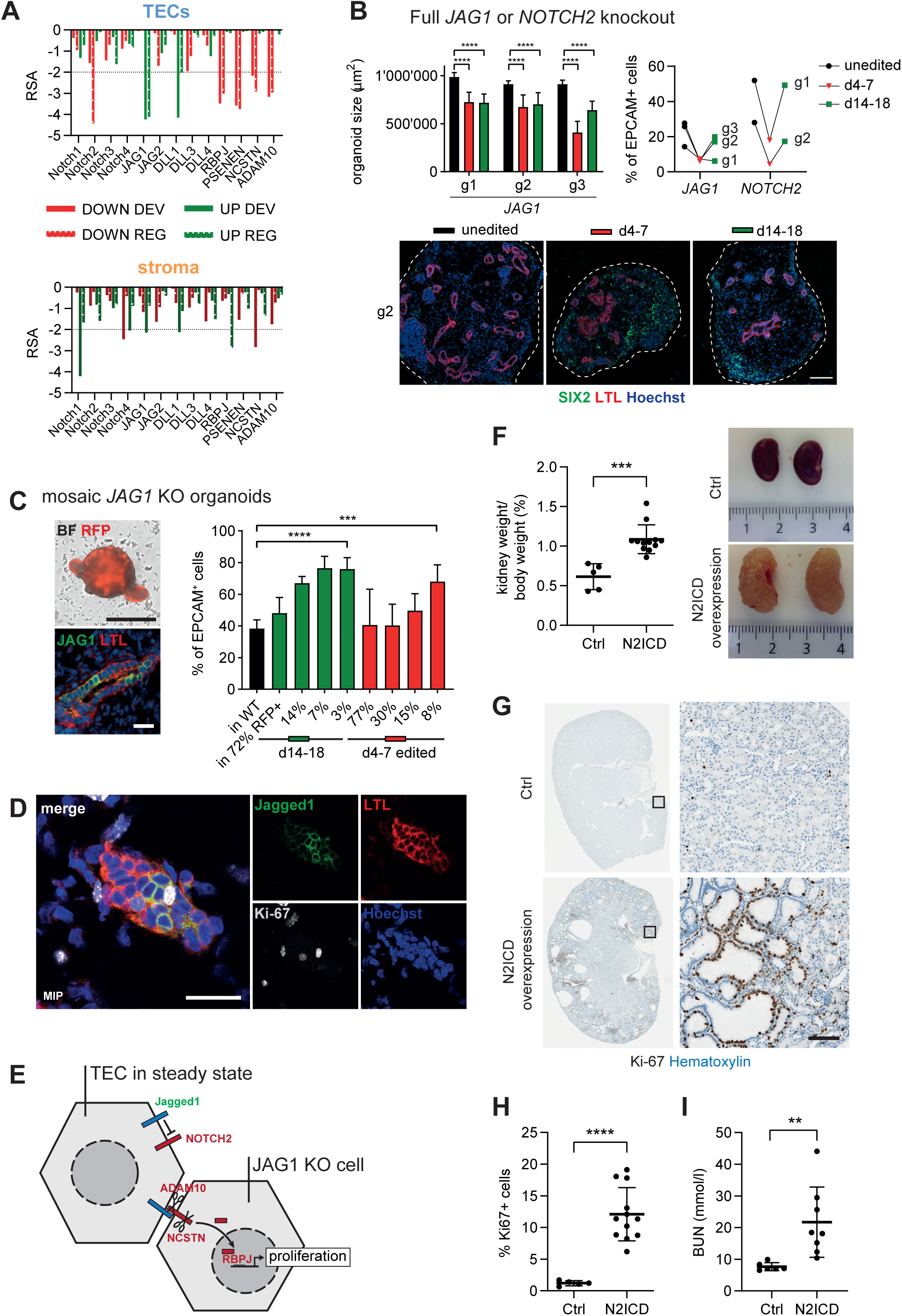
A novel cis-inhibitory effect of Jag1 in kidney tubular epithelial cells. (A) RSA scores for Notch signaling genes in DEV and REG axis for depletion (DOWN) or enrichment (UP) separated by cell type. (B) KO of *JAG1* or *NOTCH2* in organoids. iPSCs were infected with lentiviruses delivering 3 different sgRNAs (g1, g2, g3) targeting *JAG1* or *NOTCH2* and differentiated into kidney organoids. Organoid size was significantly (*p* < 0.0001, unpaired two-tailed Student’s t-test) decreased after *JAG1* KO induced between day 4-7 or 14-18 (n = 9-12 organoids each). At the same time, the number of EPCAM^+^ cells, assessed by flow cytometry at day 35, dropped upon *NOTCH2* or *JAG1* depletion. Representative whole organoid sections immunostained for proximal tubular cells (LTL) and the NPC marker SIX2 corroborates a defect in nephrogenesis upon *JAG1* depletion. Scale bar 100 µm. (C) Mimicking the pooled nature of the CRISPR screen, mosaic cultures between WT and *JAG1* gRNA containing cells (RFP^+^) were differentiated into organoids. Brightfield/RFP overlay of an organoid initiated with ¼ *JAG1* g2 cells illustrates the sporadic growth of *JAG1* KO cysts (RFP^+^). Scale bar 1 mm. Immunostained proximal tubule illustrates the mixed nature of ¼ JAG1 organoids. Scale bar 20 µm. At day 35, the number of EPCAM^+^ cells in the final RFP^+^ cell population (potential *JAG1* KO) was quantified by flow cytometry and revealed a significant overrepresentation of tubular cells in the *JAG1* KO fraction. The tubular fraction increases the more the percentage of *JAG1* KO cells (n = 7-8 organoids each) gets diluted in WT cells (n = 6 organoids) (*** *p* = 0.0002, **** *p* < 0.0001, unpaired two-tailed Student’s t-test). Mean ± s.d.. (D) Immunofluorescence stainings on cryosections from day 35 organoids initiated with ½ *JAG1* g2 cells suggests that Jagged1 expressing WT cells may induce proliferation (Ki-67) in neighboring Jagged1^-^ tubular cells. Scale bar 20 µm. (E) Cartoon illustrating the effect of Jagged1 on Notch signaling in cis and in trans. Upon complete *JAG1* KO no positive signal in trans can induce proliferation, in agreement with the necessity of NOTCH2, RBPJ, NCSTN and ADAM10 for proliferation within the tubular lineage. In case of mosaic depletion of Jagged1 (as in a pooled CRISPR screen), the *JAG1* KO cells loses its cis-inhibition and is more responsive for signaling in trans. (F) Kidneys are increased in size in N2ICD overexpressing mice (n = 12) 11 months after TAM injection compared with WT (n = 5) littermates (*** *p* = 0.0002, unpaired two-tailed Student’s t-test). The % kidney weight was calculated using the average weight from both kidneys. Mean ± s.d.. (G) Histopathology reveals multiple, large fluid-filled cysts in the cortex and medulla containing cycling tubular epithelial cells (Ki-67^+^). Scale bar 100 µm. (H) Percentage of Ki-67 positive cells in N2ICD (n = 12) overexpressing TEC mouse kidneys increases significantly compared to WT kidneys (n = 5) (**** *p* < 0.0001, unpaired two-tailed Student’s t-test). Ki-67 positive cells were quantified by an automated nuclear count algorithm in Imagescope on Ki- 67/Hematoxylin IHC whole kidney sections shown in (G). Mean ± s.d.. (I) Blood urea nitrogen levels are increases in N2ICD mice (n = 8) 11 months after tam injection compared with WT (n = 6) littermates (** *p* = 0.0088, unpaired two-tailed Student’s t-test). Mean ± s.d..

To investigate the role of Jagged1 in cis-inhibition (Bray, 2016), we generated iCas9 iPSC lines containing different sgRNAs targeting *JAG1* or *NOTCH2*, differentiated the lines and induced editing either before tubular specification (day 4-7) or after lineage choice (day 14-18). All *JAG1* gRNAs led to reduced organoid growth and a reduction in the number of epithelial tubules in both editing regimes (Figure 7B). We observed the same effect upon *NOTCH2* KO, suggesting that Jagged1 is the ligand for Notch2 in promoting tubular specification. Further, we show an arrest of cells at the nephron progenitor stage upon early *JAG1* KO (SIX2^+^ cells in mid panel in Figure 7B) consistent with a role of Notch signaling in regulating NPC self-renewal versus differentiation (Mukherjee et al., 2019).

The necessity of Jagged1 for the tubular lineage does however not explain how a *JAG1* KO resulted in more EPCAM^+^ cells in our screen. To mimic the mosaic KO of *JAG1* as in the pooled screen, we performed a dilution series of polyclonal iCas9 gRNA_JAG1 cells with WT cells. The final number of potential *JAG1^-/-^* cells in the organoid was assessed at the endpoint by RFP fluorescence (expressed by the gRNA lentivirus) with flow cytometry and ranged from 72% to 3%. The lower the ratio of *JAG1*^-/-^ to WT cells, the stronger they enriched in the tubular fraction (Figure 7C and Figure S6C). Further, already a 1:2 mixed organoid grew normal in size and we observed some small RFP^+^ cysts (Figure 7C and Figure S6B). Strikingly, we found cycling tubular cells that were Jagged1-negative in contact with Jagged1-expressing cells in mature organoids (Figure 7D). These data support a model in which Jagged1 also counters the inductive, pro-proliferative signal by inhibiting Notch2 in *cis* (Figure 7E). In simple terms, the balance between ligand and receptor will initially (in the presence of more Notch2) induce tubulogenesis and at later stages (with more Jagged1 present) stop the inductive cue.

To assess the importance of this cis-inhibitory control by Jagged1 on Notch2 *in vivo*, we generated an inducible mouse model overexpressing the intracellular domain of Notch2 (N2ICD) in epithelial cells (N2ICD het x HNF1β-CreERT2 wt), allowing us to overcome the inhibitory signal of Jagged1. Induced N2ICD overexpression in adult kidney tubule epithelium led to a significant increase in kidney to body weight ration and blood urea nitrogen 11 month after tamoxifen injection (Figure 7F&I). Histological analysis of the kidney showed renal cysts in the cortex and medulla and increased proliferation in cyst lining epithelial cells (Figure 7G&H, Figure S6D). These data strongly support that cis-inhibition by Jagged1 regulates tubular proliferation even in an adult setting. The necessity of tight control of Notch2 signaling in kidney tubules becomes also apparent in Hajdu-Cheney syndrome (Mukherjee et al., 2019). Patients with truncating mutations in Notch2 that affect the PEST domain and thus lead to a longer half-life of N2ICD display cystic kidneys – the same patho-phenotype we observe *in vitro* upon mosaic Jagged1 depletion and in mice upon N2ICD overexpression.

## Discussion

Here we described the first genome-wide pooled CRISPR screen in kidney organoids which generated a large dataset of genes functionally contributing to human nephron development resolved over time and cell types. Compared to pooled genetic screening in organoids of tumor models which rely on clonal outgrowth (Michels et al., 2020; Ringel et al., 2020), we have demonstrated that functional genetic screening is possible while the complex 3D structure of organoids is kept intact to fully leverage its potential of dissecting cellular processes which depend on the immediate environment and cell-cell contacts. For this benchmark study, we relied on a fully chemically defined iPSC differentiation protocol (Morizane and Bonventre, 2017) and focused our efforts on the stromal and tubular compartment. We expect other kidney organoid protocols which give rise to even more comprehensive kidney structures, including e.g. collecting duct or vascularization (reviewed in Nishinakamura (2019)), to be suitable for whole-genome screening, too, as long as upscaling allows production of sufficient cell culture mass.

A dox-inducible Cas9-system has proven itself very useful, because it allowed us to induce gene editing at the iPSC level (DEV) and after the last lineage choice into tubular, stromal or podocyte cell types (REG), respectively. Notably, the spatio-temporal induction of Cas9 has to be chosen carefully to take place still early enough allowing sufficient cell duplications to happen which only then will eventually lead to a quantifiable readout in barcode sequencing. Besides evaluating which genes are important for the induction of the metanephric mesenchyme, KO at iPSC level (DEV) can potentially also be informative about which genes play a role in the contribution to later cell fates (e.g. nephron structures or TECs). However, gene editing in iPSCs does not allow to dissect at which level throughout the whole differentiation process individual genes act. This holds also true for genes which act during multiple steps of nephrogenesis (e.g. Notch signaling) demonstrating that KO after day 14 of the differentiation protocol (REG) is a critical element of this study. For example, Notch signaling is known to play an important role during nephron patterning and was suggested to be involved in kidney regeneration (Mukherjee et al., 2019). We cover multiple roles of Notch signaling in our screen (exit from NPC stage and tubular specification). Moreover, we were able to dissect out a novel cis-inhibitory mechanism of Jagged1 in late tubular proliferation preventing cystic overgrowth of TECs in adult nephrons which we confirmed in an *in vivo* model.

Our screen serves as a valuable resource for the kidney research community, as it nominates genes which upon LoF may cause kidney patho-phenotypes in humans. We show here on two examples, novel CAKUT genes and ciliopathies, that our screen captures disease genes and provides a first impression on the expected phenotype. With this, we suggest that our dataset can serve as prioritization engine for LoF patient variants from WES. Nonetheless, there is still much to explore in our dataset. Many genes which we identify as hits likely contributing to kidney organoid development remain uncharacterized (e.g. the STAGA complex) or have no association with any gene set. Of the 161 genes affecting TECs at a late developmental stage, 74 cannot be assigned to any gene set in GSEA. While hit scoring and follow-up of known factors confirms the validity of our approach (e.g. ROCKi), these completely novel functional associations of genes to human kidney development make up the strength of a genome-wide genetic screen and will increase our systematic knowledge of complex biological processes. Our dataset will thus provide the kidney field with an important systems-scale overview for hypothesis generation and serve as a proof-of-concept study allowing future genetic screens both in kidney organoid disease models and in organoids of other tissues.

## Acknowledgments

We would like to thank Matthias Müller for sharing the iPSC lines used in this paper and Lapo Morelli for technical assistance. We would also like to thank Tewis Bouwmeester, Piérre Saint-Mézard, Lorna Hale, Georgios Kalamakis and Lukas Bammert for critically commenting on the manuscript.

## Author contributions

R.U., G.R., J.T., D.H. and P.S.H. designed and supervised experiments. R.U., L.G. and M.-C.L. coordinated, optimized and performed iPSC and organoid cultures. A.W. and R.C. performed RNA sequencing experiments. A.B. provided and advised on collagen-fiber staining. R.U., L.G., M.-C.L. and P.S.H. performed the CRISPR screen. W.C. performed NGS sequencing of the CRISPR screen. R.U., L.G. and M.-C.L. performed organoid follow-up experiments. J.T. provided mouse lines. V.O. performed mouse experiments. M.B. analyzed RNA-seq data. F.N. provided the pipeline and advised on CRISPR screen analysis. R.U. analyzed the CRISPR screen and performed pathway analyses. R.U. and F.N. performed statistical analysis. R.U. and P.S.H. wrote the paper.

## Declaration of interests

All authors are or were employees of Novartis Pharma AG and may own stock in the company.

## Supplemental Figure Legends

**Figure S1.**
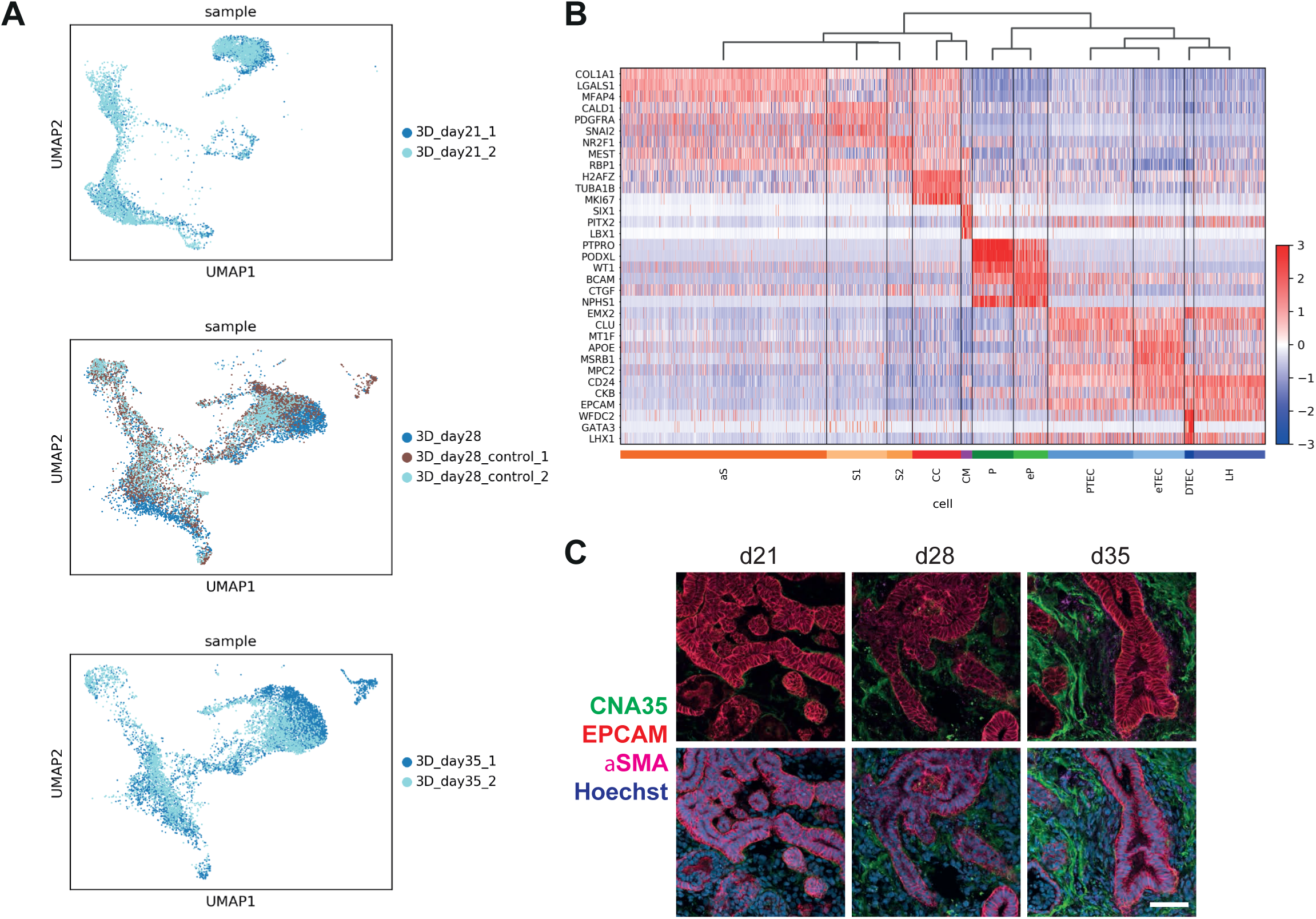
scRNA-seq QC and heatmap with marker genes (related to Figure 1) (A) UMAP of differentiation replicates at day 21, 28 and 35. (B) Expression of 3 specific marker genes for each cluster (see Table S1). Cell types are hierarchically clustered and gene expression is row normalized. (C) Immunostaining on cryosections for collagen fibers (CNA35), tubular cells (EPCAM) and activated stroma (*α*SMA). Scale bar 50 µm.

**Figure S2.**
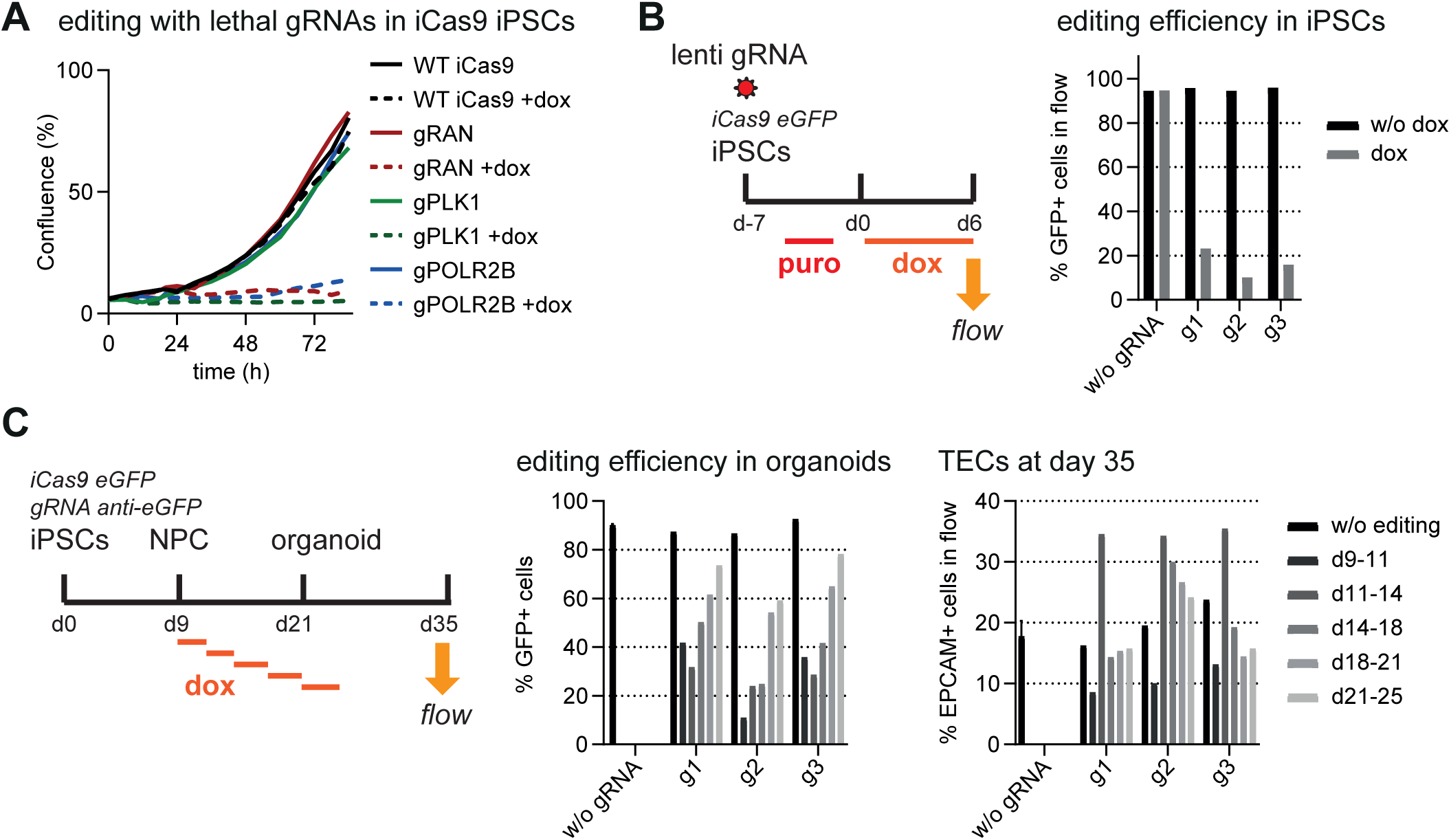
Tightness of inducible genome editing system and frequency of indel generation in iPSCs and during kidney differentiation (related to Figure 1) (A) Inducible genome editing targeting lethal genes affirms tight control over Cas9 expression. (B) GFP editing in iPSC shows efficient indel generation upon dox, quantified by flow cytometry. (C) GFP editing on organoids at different time intervals during differentiation. Note that editing on organoids at d9-11 or d11-14 influences epithelial development.

**Figure S3.**
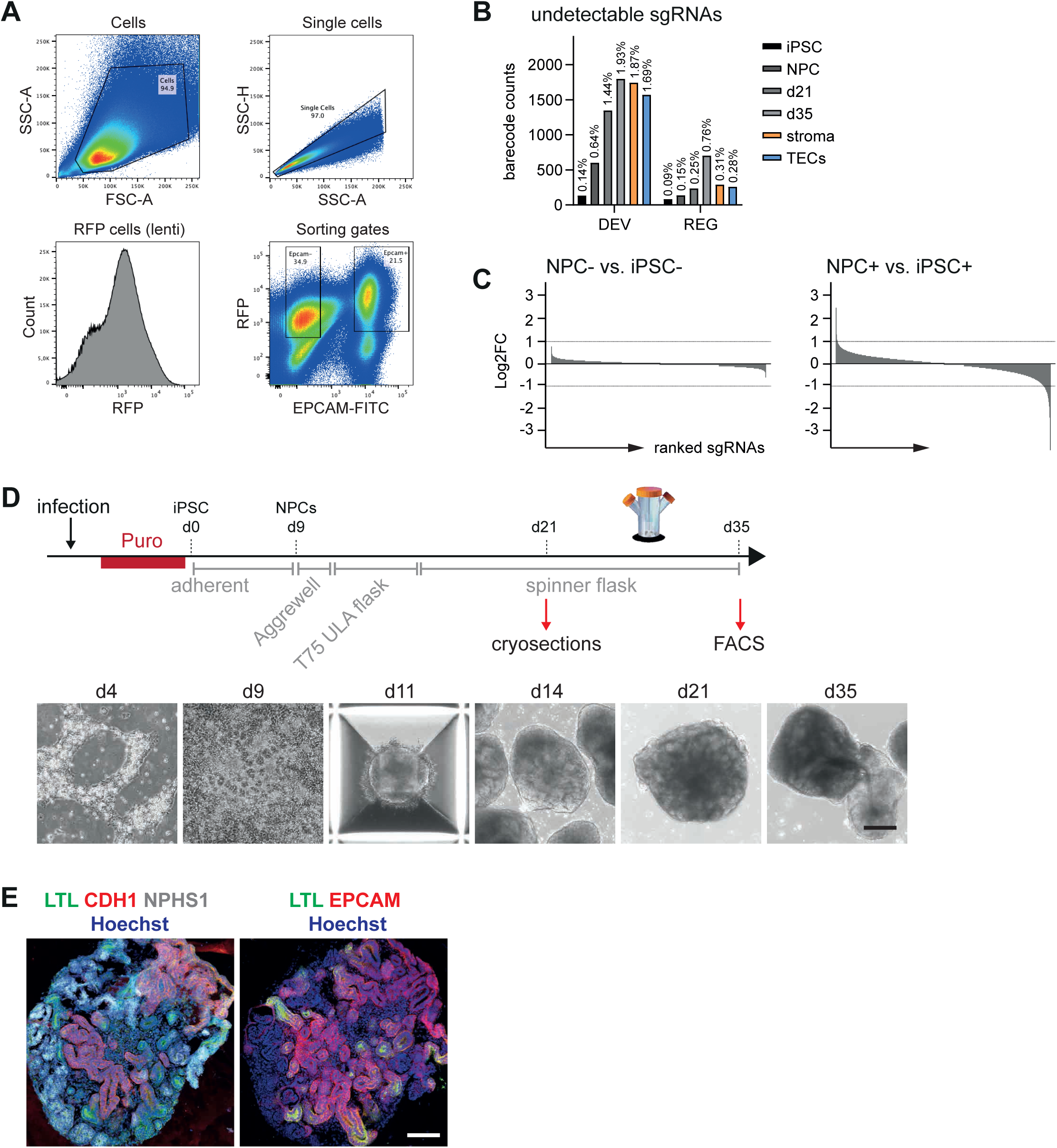
CRISPR screen QC (related to Figure 2) (A) Sorting regime for CRISPR screen. Fixed cells were stained with EPCAM-FITC and sorted for purity to obtain an EPCAM^+^ tubular cell fraction and an EPCAM- ‘stromal’ cell fraction (coverage of min. 300 cells per sgRNA). Note that EPCAM intermediate cells are podocytes that were not collected. (B) Bar plot showing absolute numbers of missing barcodes (sgRNAs) in individual samples. The numbers above the bars represent percentage of undetectable gRNAs from the whole library. (C) Waterfall plots illustrate the lack of change in barcode count in NPCs without editing in contrast to increased fold changes in NPCs with editing. With a coverage of 300 cells per sgRNA we successfully prevented clonal effects/bottlenecks within the first 9 days of differentiation. (D) Scheme illustrating upscaling of organoid cultures for genome-wide screening. Brightfield imaging over the course of differentiation. Scale bar 100 µm. (E) Cryosections and immunostaining at day 21 of culture shows that organoids in screening setup developed segmented tubules, podocytes and connective tissue. Scale bar 100 µm.

**Figure S4.**
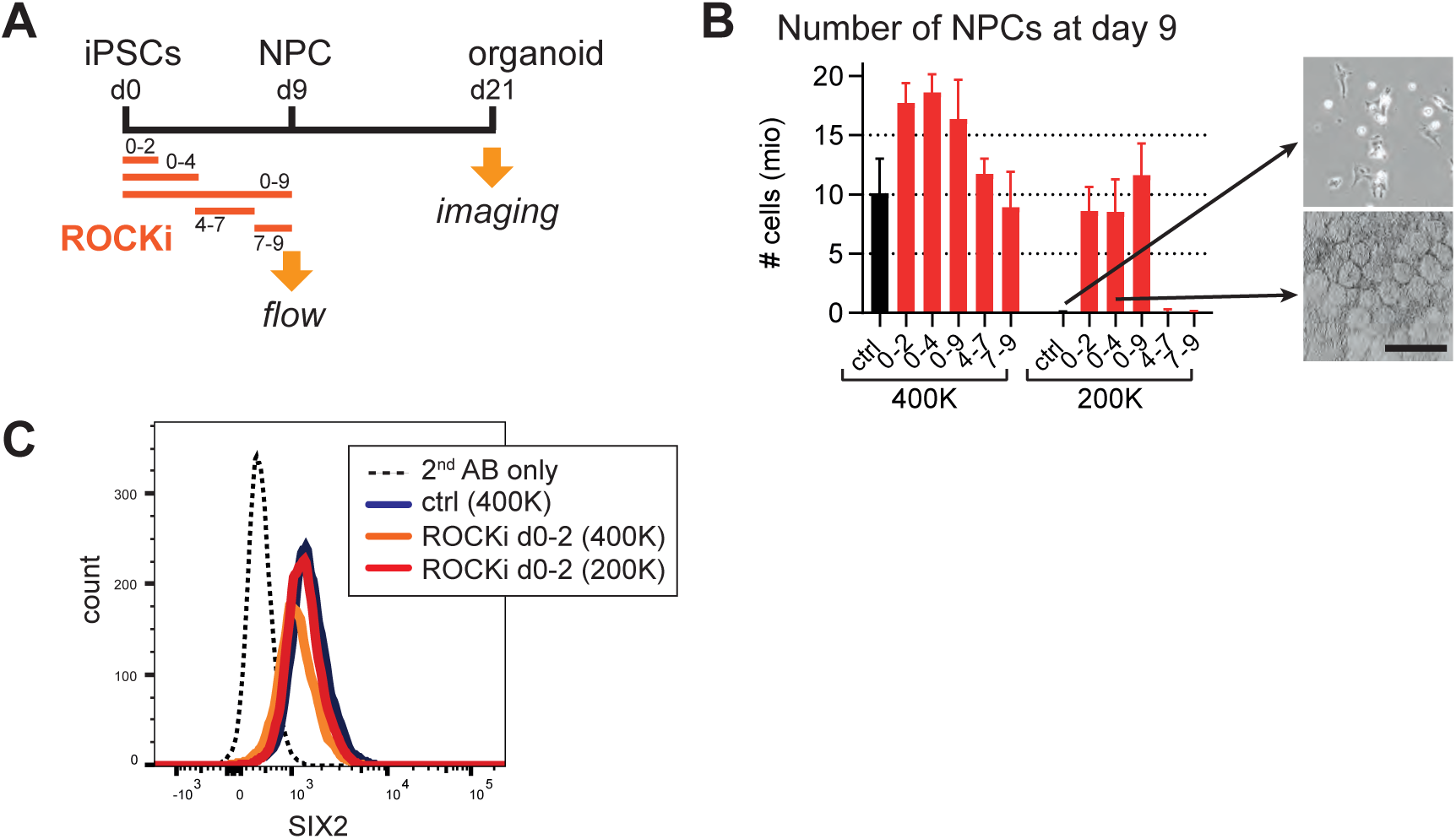
Timing of ROCKi treatment during kidney organoid differentiation (related to Figure 3) (A) Experimental setup for testing the effect of chemical ROCK inhibitor in early specification towards NPCs. (B) Treatment of cells with ROCKi during the first 2 days of differentiation leads to pronounced increased survival, as assessed by absolute cell number at day 9 of differentiation. Effect is more apparent at lower initial seeding density (200K: 200000 cells seeded per 6-well). N=2. Representative brightfield images illustrate successful induction and initial 3D structuring of NPCs at day 9. Scale bar 100 µm. (C) SIX2^+^ NPCs at day 9 quantified by flow cytometry. All conditions in (D) show overlapping histograms for SIX2. Minimal treatment of 2 days necessary for an effect is shown.

**Figure S5.**
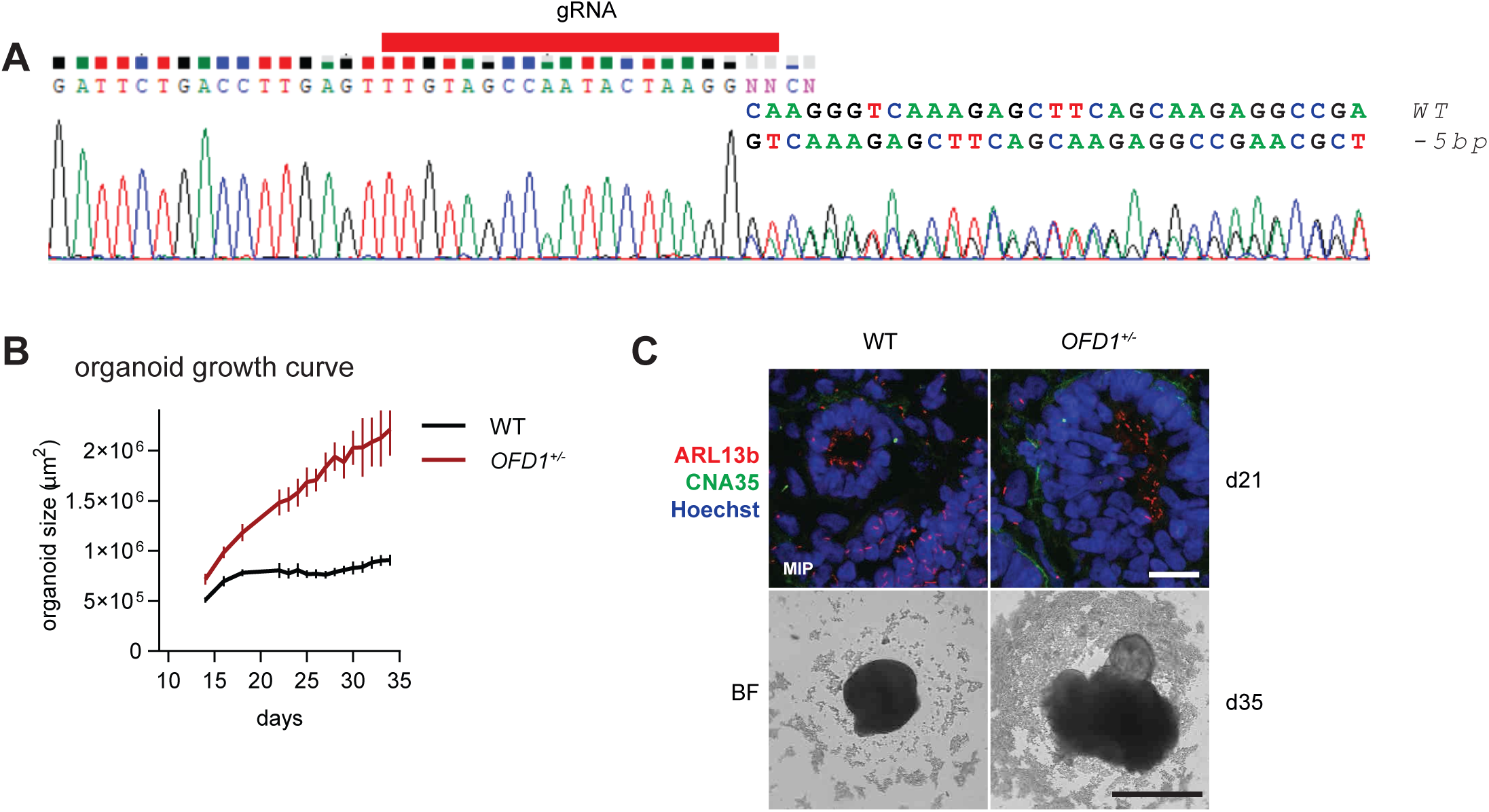
OFD1 heterozygous clone (related to Figure 5) (A) Sanger sequencing validation of sequence changes introduced by synthetic gRNA targeting OFD1 exon 16 (hg38 chrX:13,734,748-13,769,357). Sequence decomposition revealed a 5 bp deletion and a WT allele. (B) Longitudinal tracking of organoid size by brightfield imaging (n=16). (C) Immunofluorescence staining on cryosections from day 21 organoids shows the presence of cilia (ARL13b) in OFD1^+/-^ tubular epithelial and non-epithelial cells. Scale bar 50 µm. Size increase and cystic growth visible in brightfield images at the endpoint. Scale bar 1 mm.

**Figure S6.**
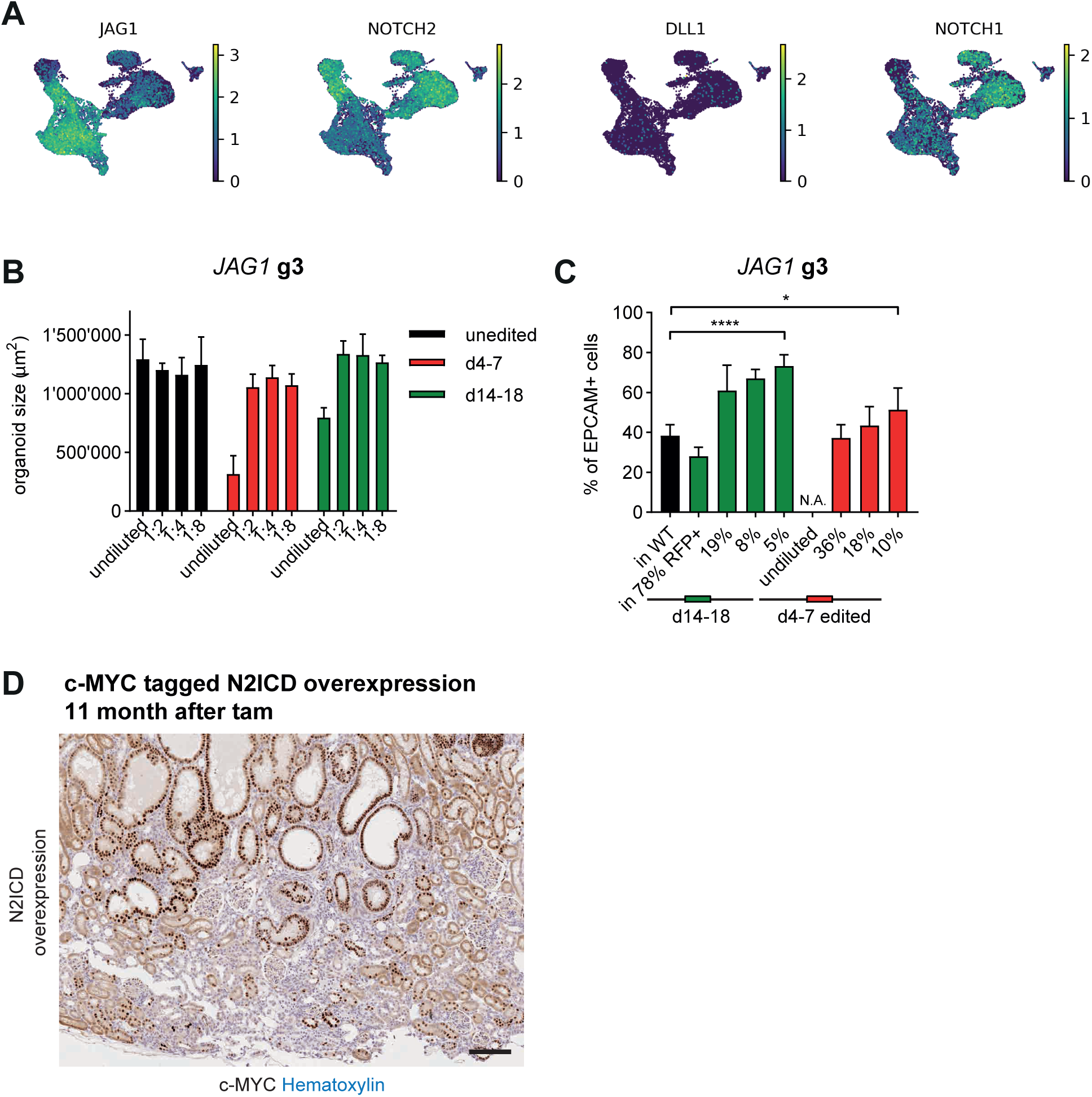
Notch signaling components in *in vitro* human and *in vivo* kidney development (related to Figure 7) (A) Expression of Notch signaling components in scRNA-seq data. Feature plot UMAP displays expression of JAG1, NOTCH2, DLL1 and NOTCH1. Note the strong expression of JAG1 in tubules, whereas NOTCH2 is expressed 2-fold higher in stroma and podocytes compared to tubules. The NOTCH1 receptor is predominantly expressed in stromal cells, consistent with an effect of NOTCH1 deletion on the stroma in the screen. (B) Organoid size at day 35 for undiluted and mosaic organoids between iCas9 empty and *JAG1* g3 expressing cells. Note that already in a 1:2 mixed organoid the size is normal, while in an undiluted setting the *JAG1* KO leads to very small organoids and misdevelopment. (C) Same experiment as in Figure 7C for *JAG1* g3: At day 35, the number of EPCAM^+^ cells in the final RFP^+^ cell population (potential *JAG1* KO) was quantified by flow cytometry and revealed a significant overrepresentation of tubular cells in the *JAG1* KO fraction. The tubular fraction increases the more the percentage of *JAG1* KO cells (n = 8 organoids each) gets diluted in WT cells (n = 6 organoids) (* *p* = 0.0218, **** *p* < 0.0001, unpaired two-tailed Student’s t-test). Mean ± s.d.. (D) Histopathology in c-MYC tagged N2ICD het HNF1β-CreERT2 mice at 11 months after tam injection. Note the high number of c-MYC^+^ nuclei in dilated tubules. Scale bar 100 µm.

## METHODS

### RESOURCE AVAILABILITY

#### Lead Contact

Further information and requests for resources and reagents should be directed to and will be fulfilled by the Lead Contact, Philipp Hoppe

#### Materials Availability

There are restrictions to the availability of the whole-genome library due to Novartis Pharma AG policies. All other unique/stable reagents generated in this study are available from the Lead Contact with a completed Materials Transfer Agreement.

#### Data and Code Availability

RNA-sequencing data of the organoids will be deposited to a public repository. Original/source date will be made available on Mendeley Data. The de-identified barcode counts, barcode fold-changes, barcode–gene associations and gene results generated during this study can be made available as supporting information. Code for RSA analysis and Jupyter Notebooks are available upon request.

### EXPERIMENTAL MODEL AND SUBJECT DETAILS

#### Human induced pluripotent stem cells

For all experiments where no other donor cell line is indicated, wt29 female human induced pluripotent stem cells (derived from cell line AG09429 from the NIA Aging Cell Repository at the Coriell Institute for Medical Research) were used. Donors 2 and 3 (Figures 3C-D) have been described previously (Bidinosti et al., 2016). iPSCs were cultured in mTeSR (Stem Cell Technologies) supplemented with Penicilin/Streptomycin (Gibco) on laminin-521 (Biolamina) coated plates. For routine maintenance, approx. 70% confluent cultures were dissociated into single cells with TrypLE (Gibco) and seeded at a seeding density of 10000-25000 cells/cm^2^ in mTeSR supplemented with 2 µM ROCK inhibitor Y27632 (Tocris) onto laminin-521 coated plates. After 24h, the medium was replaced with mTeSR without ROCKi followed by daily medium changes.

Inducible Cas9 iPSCs were generated using an AAVS1 TALE-Nuclease KIT (GE60xA-1; System Biosciences) and an iCas9 HR donor plasmid containing a TRE3G promoter-driven spCas9 with a C-terminal MYC NLS followed by a CMV promoter-driven Tet-On-3G and a neomycin resistance adapted from Ihry et al. (2018). The plasmids were delivered at equal ratio by lipofection with Lipofectamin Stem (Invitrogen). After G418 selection (100 μg/ml; Gibco), clones were picked, expanded and screened by treating with doxycycline (500 ng/ml; Clontech) for 40 h and staining for Cas9 (7A9-3A3, Cell Signaling) by immunofluorescence. Clones with homogenous expression of Cas9 and tight control in the absence of dox were subsequently banked and tested for proper targeting using junction PCR. The final clone used for genome-wide screening contained a single, heterozygous insertion of iCas9 at the AAV1 locus. Karyotyping was performed for WT and iCas9 iPSC lines and showed no clonal abnormalities.

#### Mouse strains

Mice allowing Tamoxifen (TAM)-inducible expression of Cre (CreERT2) in HNF1β^+^ kidney tubular epithelial cells were generated by targeting an ires-CreERT2 cassette to the 3′-UTR of the mouse Hnf1β gene locus in C57Bl/6 ES cells. Successfully targeted ES cells were injected into pseudopregnant foster mice and chimeric offspring was mated with C57Bl/6 mice to confirm germline transmission, yielding B6-HNF1β^tm1(CreERT2)Npa^ mice; termed HNF1β-CreERT2 thereafter). As a next step these mice were crossed with mice allowing for Cre-inducible expression of the Notch2 intracellular domain (N2ICD mice, (Tchorz et al., 2009)), yielding N2ICD/HNF1β-CreERT2 mice. To induce N2ICD expression in HNF1β^+^ cells, and as a control (ctrl), 8 - 13 weeks old mice were injected i.p. with 100 µl of TAM (20 mg/ml in sunflower oil) once daily for 5 consecutive days. Mice were analyzed 11 months after induction. 5 ctrl (N2ICD^het^/HNF1β-CreERT2^-^) (3 female (f), 2 male (m)) and 12 N2ICD (N2ICD^het^/HNF1β-CreERT2^het^) (5 f, 7 m) mice were used for Figure 7F-H, and 6 Ctrl (2 f, 4 m) and 8 N2ICD (3 f, 5 m) mice were used for Figure 7I. Bodyweight at the beginning of the study was 24,7 g (mean) ± 1,4 g (s.d.) for WT f mice, 23,4 g ± 1,8 g for N2ICD f mice, 31,3 g ± 2,1 g for WT m mice and 31,4 g ± 2,0 g for N2CID m mice.

Animals were monitored daily for wellbeing. All mice had unrestricted access to water and food and animal experimentation was conducted in accordance with animal law of Basel-Stadt, Switzerland. The Novartis Campus animal facilities comprise an SPF animal breeding facility and a clean facility for experimental surgery and physiology. Biosecurity and pathogen exclusions follow the Federation of European Laboratory Animal Science Associations (FELASA) health monitoring guidelines, and animals are screened quarterly. Mice were housed in individually ventilated cages in a 12:12 light:dark cycle. Environmental enrichments included nest-lets, wood sticks and mouse houses. The maximum housing density in the breeding and experimental areas was 4 and 5 mice, respectively. Most experiments were performed with grouped mice, single housing was used for aggressive or wounded animals.

### METHOD DETAILS

#### Kidney organoid differentiation

iPSCs were dissociated into single cells with TrypLE (Gibco) and plated onto laminin-521 coated plates at a seeding density of ∼54000 cells/cm^2^. The plated cells were cultures in mTeSR supplemented with 2 µM Y27632 for 8-16h before starting differentiation. To induce primitive steak, cells were washed with PBS and cultured for 4 days in basic differentiation medium (Advanced RPMI 1640, 1x GlutaMAX, Pen/Strep; all Gibco) supplemented with 8 µM CHIR99021 (Tocris) + 5 ng/ml human Noggin (Peprotech) with a medium change after 2 days. At day 4 the cells were carefully monitored to see clustering of cells with a shiny border, a characteristic of the right time point to change to differentiation into posterior intermediate mesoderm. For this, cells were incubated for 3 days in basic differentiation medium containing 10 ng/ml activin A (R&D). Next, nephron progenitors were induced in basic differentiation medium supplemented with 10 ng/ml human recombinant FGF9 (R&D). At day 9, nephron progenitor cells (NPCs) can be dissociated into single cells with TrypLE, frozen for later culturing in Cryostore CS10 (Stem Cell Technologies) at a cell density of 5 mio/ml or aggregated into 3D suspension culture. For this 10000-100000 NPCs were seeded in basic differentiation medium with 10 ng/ml FGF9, 3 µM CHIR99021 and 2 µM Y27632 into 96-well ULA round bottom (Corning) or Aggrewell800 plates (Stem Cell Technologies). To induce aggregation, plates were spun for 3 min at 90g. The two consecutive days, half of the medium was replaced with basic differentiation medium supplemented with 10 ng/ml FGF9. From day 14 onwards, organoids were cultured in basic differentiation medium without any growth factors and half of the medium was changed every 2 days.

#### Organoid single cell dissociation

For flow cytometry, organoids were dissociated with a 1:1 mix of 0.25% Trypsin-EDTA and non-enzymatic cell dissociation solution (both Gibco) for 20 min at 37°C. After 10 and 20 min, organoids were broken up by pipetting (up and down 20x). Then cells were washed twice with 10% and 1% FBS in PBS. For scRNA sequencing the cell suspension was additionally passed through a 40 µm and a 20 µm filter.

#### Single cell RNA sequencing and data processing

The single cell organoid cell suspension was resuspended in PBS with 0.05% BSA. Viability was accessed by Trypan blue cell counting and exceeded 90%. Cells (2000–10,000) were loaded on a 10× Genomics Chromium instrument to generate single-cell GEMs. scRNA-Seq libraries were prepared using a GemCode Single Cell 3′ Gel Bead Kit according to the manufacturer’s instructions (10× Genomics). Amplified cDNA products were cleaned up with the SPRIselect Reagent Kit (0.6× SPRI). Indexed sequencing libraries were constructed using the reagents in the Chromium Single Cell 3′ library kit V2 (10× Genomics). The barcoded sequencing libraries were quantified using a Qubit 2.0 dsDNA HS Assay Kit (Invitrogen). The quality of the libraries was accessed on an Agilent 2100 Bioanalyzer using an Agilent High Sensitivity DNA kit. Sequencing libraries were loaded at 10 to 12 pM on an Illumina HiSeq 2500 platform.

Expression data for 24’781 cells and 33’694 Ensembl genes obtained from cellranger (10X) were processed with a generic workflow based on scanpy version 1.4.4.post1 as follows. First, a series of quality filters to exclude low quality cells was applied. Five cells that expressed less than 724 genes and 9’826 genes that were present in fewer than three cells were excluded. In addition, cells which expressed more than 5000 genes were excluded as potential doublets. Further we required a percentage of mitochondrial reads of less than 15% per cell. Finally, all cells with less than 1449 UMIs were also excluded. This resulted in 20’055 high quality cells. Expression values for these cells were normalized per cell and log transformed. We then determined highly variable genes (min_mean=0.01, max_mean=3, min_disp=0.5). On the set of highly variable genes expression values were normalized by regression on the total number of UMIs per cell and the percentage of mitochondrial reads per cell and finally scaled to a maximum value of 10. We then computed the neighborhood graph using 20 neighbors and 40 principal components. Based on this neighborhood graph a UMAP representation and several Louvain clusterings (resolutions: 1, 0.5, 0.25 and 0.05) were obtained. In order to annotate the different clusters, marker genes were determined using Wilcoxon tests. As a means to check for the influence of cell cycle status we used the score_genes_cell_cycle function of scanpy together with a commonly used list of marker genes (https://github.com/theislab/single-cell-tutorial/blob/master/Macosko_cell_cycle_genes.txt).

#### Immunofluorescence and organoid imaging

Live organoid imaging was performed at an Incucyte S3 using the spheroid module for autofocusing and image analysis (Sartorius).

For higher resolution imaging, organoids were fixed in 4% PFA in PBS for 30 min at room temperature (RT) followed by overnight incubation in 50% sucrose. Next, organoids were embedded in a small block of 7.5% gelatin (from porcine skin, Sigma) and 10% sucrose. After solidification of the gelatin block for 6h at 4°C it was embedded in a larger OCT block (Qpath, VWR) and subjected to cryosectioning (10 µm) using a Leica Cryostat.

Kidney organoid cryosections were washed in PBS for 10 min to remove residual OCT followed by permeabilization (if necessary) and blocking in 0.2% Triton X-100 in 2%BSA/PBS for 30 min at RT. For Lotus Tetragonolobus Lectin (LTL biotinylated) staining sections were additionally blocked with a streptavidin/biotin blocking kit (both Vector Laboratories) according to the manufacturer’s instructions. Subsequently, sections were incubated with primary antibodies for 1h at RT or overnight at 4°C. Sections were then washed three times with PBS/0.4% BSA followed by 1h incubation with secondary antibodies and Hoechst at RT. Sections were again washed three times with PBS/0.4% BSA and mounted with ProLong Diamond antifade mountant (Invitrogen). Confocal images were captured using a Zeiss LSM900 laser-scanning microscope (Carl Zeiss) equipped with 20x, 40x or 63x oil-immersion objectives and an Airy scan. Images were visualized using ImageJ software.

The following antibodies/staining reagents were used in this study: mouse polyclonal anti-ARL13b (Proteintech), mouse monoclonal anti-aSMA (DAKO), mouse monoclonal anti-Cas9 (Cell signaling), rabbit monoclonal anti-CDH1 (Thermo Fisher), mouse monoclonal anti-EPCAM-AF647, mouse monoclonal anti-EPCAM-FITC, rabbit monoclonal anti-JAG1 (all Abcam), rat monoclonal anti-Ki67 (eBioscience), sheep polyclonal anti-NPHS1, mouse monoclonal anti-PODXL (both R&D System), rabbit polyclonal anti-SIX2 (Proteintech), donkey anti-Sheep IgG (H+L) antibody Alexa Fluor® 488, donkey anti-Mouse IgG (H+L) antibody Alexa Fluor® 488, donkey anti-Mouse IgG (H+L) antibody Alexa Fluor® 594, donkey anti-Rat IgG (H+L) antibody Alexa Fluor® 594, donkey anti-Rabbit IgG (H+L) antibody Alexa Fluor® 488, donkey anti-Rabbit IgG (H+L) antibody Alexa Fluor® 594 (all Thermo Fisher), Lotus Tetragonolobus Lectin (LTL) Biotinylated (Vectorlab), Fluorescent Dye 633-I Streptavidin (Abnova), Hoechst (Thermo Fisher).

#### CNA35 production, labelling and staining

CNA35-Alexa488 was produced and labelled internally. The DNA encoding for domains N1 and N2 of the A-region of Staphylococcus aureus collagen adhesion protein (CNA35, amino acids 30-344) optimized for expression in *E. coli* and with flanking NdeI (5’) and XhoI (3’) restriction sites was ordered from GeneArt and cloned into the pET26b (Novage) expression vector. QuickChange (Stratagene) was used for Y175K mutagenesis and LIC-PCR (Lifetechnologies) was applied for the introduction of ggc-tct-tcc-ctg-ccg-gaa-acc-ggt-ggc (encoding GSS-LPET/G-G, GSS-SrtA-tag) at the 3’ end of the CNA35 sequence, to allow site-specific labeling of CNA35 using recombinant sortase A. Each of the three resulting plasmids (MRGS-His6-CNA35, MRGS-His6-CNA35-SrtA, MRGS-His6-CNA35(Y175K)) was transformed into chemically competent *E. coli* BL21 (DE3) Gold cells (Stratagene). For large scale expression, 50 mL preculture (LB medium: 0.5 g/L NaCl (Merck), 10 g/L tryptone (Bacto), 5 g/L yeast extract (Bacto), 6 g/L glucose (Roquette), 25 g/L kanamycin sulfate and 20 g/L tetracycline hydrochloride (both Sigma-Aldrich)) was inoculated using a glycerol stock of a preselected clone expressing CNA35, and grown at 37 °C, 260 rpm until an OD_550nm_ of 1.5 was reached. The main culture (TB medium: 0.9 L, 1.2 g/L NH4SO4 (Merck), 0.041 g/L KH2PO4 (Merck), 0.052 K2HPO4 (Merck), 12 g/L tryptone, 24 g/L yeast extract, 25 g/L kanamycin sulfate, 20 g/L tetracycline hydrochloride) was grown in batch mode in a 1 L fermenter (DASGIP 1L Bioreactor, Vaudaux Eppendorf) at pH 7.0 (+/-0.2) and 30 °C. Reduction of temperature to 25 °C and induction recombinant gene expression by addition of 1 mM IPTG (VWR) was triggered automatically by pH (7.1). Cells were harvested 5 hours after induction by centrifugation at 4000x *g*. Cells were suspended in 5 mL/g lysis buffer (50 mM sodium phosphate (Sigma-Aldrich), 500 mM NaCl (Sigma-Aldrich), 20 mM imidazole (Sigma-Aldrich), 5 mM MgCl2 (Sigma-Aldrich), 5 mM DTT (Roth), pH 8.5, supplemented with cOmplete EDTA-free protease inhibitor tablets (Roche) and Benzonase nuclease (Sigma-Aldrich) according to manufacturer’s instructions). Lysis was performed by high pressure homogenization (2 passages at 600 bar) using an APV Lab2000 homogenizer (SPX). The lysate was clarified by centrifugation and filtration, and the respective CNA35 variant was purified by IMAC (HisTrap 5ml, GE) and SEC (HiLoad 16/600 Superdex 75 pg, GE) using an ÄKTAxpress system (GE), according to manufacturer’s instructions. Labelling was performed by adding 10 % v/v of 1 M NaHCO3 solution to adjust the pH to 9. Then 6 eq. of Alexa Fluor 488 NHS ester (10 mg/mL in DMSO, Thermo Fisher) was added. The solution was incubated at RT for 4 h. The free dye was removed by purification on ÄKTA Pure (HiPrep Desalting column, PBS). The first fraction was collected and concentrated using Amicon centrifugal filter (MWCO: 10 kDa) to give CNA35-Alexa Fluor 488 randomly labeled. The product was characterized by size exclusion chromatography on a Agilent HPLC 1260 usind a Superdex 200 increase 10/300 GL column from Cytivia, and UV/Vis on a NanoDrop 1000 to determine protein and dye concentrations, and degree of labeling. To stain collagen fibers on organoid cryosections, CNA35-Alexa488 was added at a concentration determine by titration in parallel with the incubation step of secondary antibodies during immunofluorescent staining.

#### Flow cytometry

Flow cytometry was either performed on a pool of 4-12 organoids (Fig. 1F) or on individual organoids in 96- well format (dissociation in ULA round bottom plate, staining in V bottom plate) (Fig. 6D) on a BD Fortessa instrument. After organoid single cell dissociation (see above), cells were fixed and permeabilized for intracellular staining (Ki-67) using the BD cytofix/cytoperm kit according to the manufacturer’s instructions or directly stained with surface antibodies (e.g. EPCAM). Antibody incubations and washing steps were performed in PBS/0.4% BSA with centrifugation steps at 1000 (tube)-1200 (plate) rpm for 5 min at RT. 10000-100000 events were recorded on a BD Fortessa instrument.

#### Pooled CRISPR screening and analysis

We conducted a genome-scale pooled CRISPR screen covering 18360 genes (5 sgRNAs per gene whenever possible, split into two sgRNA sub-pools) (Estoppey et al., 2017). In brief, chip-based oligonucleotide synthesis was used to generate spacer-tracerRNA-encoding fragments which were PCR-amplified and cloned as a pool into the pRSI16 lentiviral plasmid (Cellecta). Lentiviral gRNA plasmids expressed a constitutive RFP and puromycin resistance to mark and select for infected cells respectively. Viral titer of the two sgRNA sub-libraries was determined by a 6-point dose response in 6-well plates by flow cytometry of RFP+ iPSCs 3 days after infection.

We desired a coverage of > 300x of each sgRNA at any time during the screen. iPSCs were infected at 0.5 MOI to ensure each cell was infected with no more than a single gRNA. This resulted in 66 mio iCas9 iPSCs cultured on 12 10 cm plates that were infected with 3.5 ml of lentivirus in total (34x dilution of HEK293 lentiviral supernatant in mTeSR). 24h after infection, iPSCs were selected with puromycin (1 µg/ml, Gibco). The next day, cells were split into 2 batches of 16 10 cm plates and cultures in either mTeSR + puro (REG axis) or mTeSR + puro + dox (200 ng/ml) (DEV axis) for 4 days with daily medium changes. At 6 days post-transduction, cells were pelleted for DNA (40 mio), banked (5 mio per cryovial) and seeded for organoid differentiation (3 mio cells per 10 cm plate; 14 plates per axis). At day 9 of differentiation, NPC were dissociated into single cells with TrypLE and counted. The DEV axis gave ∼700 mio cells, the REG axis ∼800 mio cells. 40 mio for each axis were pelleted for genomic DNA. A small portion (12-36 mio NPCs) was used for a test differentiation evaluating the nephrogenesis potential in the screening format (growth in Aggrewell plates and spinner flasks under pooled CRISPR editing conditions, see immunostainings of day 21 organoids in Fig. S3B) and final numbers achieved by FACS at the endpoint. The residual NPCs were cryopreserved in Cryostore CS10 at a concentration of 5 mio/ml. After successful QC differentiation and FACS, 288 mio NPCs per axis were aggregated into 14400 organoids (20000 cells/per organoid) in Aggrewell800 6-well plates (Stem Cell Technologies). The day after, part of the organoids were transferred to T75 ULA flasks (Corning) while medium change. The rest of the organoids was transferred at day 11 with the next medium change. At day 14, growth factors were retrieved and from day 16 onwards organoid were cultured in 500 ml spinner flasks at 100 rpm (5’400 organoids in 450 ml basic differentiation medium per flask; Corning) with regular medium changes every 2 days. Whole organoid samples for genomic DNA were collected at day 21 and 35; ∼1800 organoid per samples and axis. At day 35, ∼10’800 organoids were collected, dissociated (see *Organoid single cell dissociation*), stained with EPCAM-FITC (VU-1D9, Abcam), fixed with BD cytofix/cytoperm and sorted at a BD FACSAria Fusion for EPCAM^+^ tubular cells and EPCAM- stromal cells. Note that with restrictive gating podocytes, which are weakly EPCAM^+^ were excluded in the two sorted cell populations. For all samples, genomic DNA was isolated using phenol chloroform extraction. In short, cells were resuspended in 5 ml TNES (10mM Tris-Cl pH 8.0, 100mM NaCl, 1mM EDTA,1% SDS) and incubated overnight at 65°C to reverse PFA crosslinks. After allowing the samples to cool, samples were incubated with 100 µl RNase A (QIAGEN) for 30 min at 37 °C, followed by addition of 100 µl of proteinase K (QIAGEN) for 1 h at 45°C. 5 ml PCIA (Phenol:Cholorform:Isoamyl alcohol pH 8) (Thermo Fisher) were added and samples vortexed, spun at max speed for 2 min and the aqueous phase was transferred to 5 ml of PCIA. Samples were vortexed again, spun at max speed for 2 min the aqueous phase was transferred to 4,5 ml of chloroform. A third time, samples were vortexed, spun at max speed for 2 min and the aqueous phase was transferred to 400 µl 3M NaAcetate pH 5.2. 10 mL 100% EtOH was added, samples were mixed and DNA precipitated for 1 h on ice followed by spinning at max speed for 10 min. EtOH was decanted and pellets washed with 10 ml 70% EtOH. Finally, samples were spun at max speed for 10 min, pellets air-dried and resuspended in 1 ml nuclease free water. PCR was performed using lentiCRISPR specific primers and library construction and sequencing was performed as described by DeJesus et al. (2016).

Raw sequencing reads were aligned to the appropriate library using Bowtie (Langmead et al., 2009) allowing for no mismatches and counts were generated. Moderated fold-change estimates between samples were generated for each sgRNA using DESeq2 (Love et al., 2014). Effects on proliferation were assessed by comparison of the unsorted cell population to the input library. For gene-based hit calling, consistency of all sgRNAs per gene was considered via the RSA algorithm (König et al., 2007) and plotted against first or third quartile Z-score. RSA calculates the enrichment of all sgRNAs of a gene towards high or low ranks. Thus, RSA based ranking has the advantage that also weak gene hits can be detected by considering the effect of all sgRNAs concerned. This enables the comparison of different time points, cell types and axis in which the fold changes differ largely due to variable proliferation rates. The software Spotfire was used for data analysis and visualization.

#### Lentiviral delivery of sgRNAs for individual gene knockouts

sgRNAs (for sequences see Table S10) were cloned into the LV _TagRFP_T2A_PuroR backbone and packaged as described (DeJesus et al., 2016). iCas9 iPSCs were infected at > 0.5 MOI in a 6-well format, 8h after seeding of 200000 cells per well. After 24h, cells were maintained in puromycin (1 µg/ml), passaged once more and banked without editing. Genome editing was induced by the addition of 200 ng/ml dox. Efficiency was assessed either by protein or functional readouts (GFP fluorescence, loss in viability, ciliary ablation or JAG1 immunofluorescence staining) or locus-specific PCR and TIDE web tool analysis (Brinkman et al., 2014).

#### Synthetic sgRNA delivery by electroporation

iCas9 iPSCs were treated with dox (200 ng/ml) 24h before electroporation. Cells were detached with TrypLE (Gibco), counted using a ViCell and 100000 cells were mixed with 0.5 µg (3 µM) synthetic sgRNA (IDT) targeting OFD1 (TTGTAGCCAATACTAAGGCA). This mix was electroporated at 1200 V for 1 10 ms pulse using the Neon electroporation system (Thermo), resuspended in mTeSR and seeded onto a laminin-521 coated plate. For generation of heterozygous OFD1 clone, iPSCs were seeded by limited dilution onto a 10 cm plate. After 5 days clones were picked, expanded and screened by locus-specific PCR and TIDE web tool analysis (Brinkman et al., 2014).

### QUANTIFICATION AND STATISTICAL ANALYIS

#### Permutation analysis to estimate false discovery rate (FDR)

To estimate the significance of the RSA results they were calculated 1000 times with randomly shuffled gene–sgRNA associations. Empirical p-values were obtained as the fraction of RSA gene values that were equal or more extreme than the actual ones using the correct gene–sgRNA associations.

#### Statistical analysis

If not otherwise stated, estimation of false discovery rate (FDR) was used. Prism software (GraphPad) was used for further statistical analysis. No statistical method was used to predetermine sample size. No samples or animals were excluded from the analysis. The experiments were not randomized or blinded. Data are expected to have normal distribution. Data significance was determined by an unpaired two-tailed Student’s t-test. *n* in figure legends refers to biological replicates and is indicated.

### ADDITIONAL RESOURCES

#### Pathway analysis

GO terms, human phenotype or pathway associations were analyzed using ToppGene suite (Chen et al., 2009) or Metascape gene set enrichment (Zhou et al., 2019). Enrichment scores were calculated as log2(observed/expected) where the expected hits in case of a random overlap are calculated as ((# of hits)*(# of genes in term))/(total number of genes). The probability of finding an overlap of x genes was calculated using exact hypergeometric probability and a normal approximation (http://nemates.org/MA/progs/representation.stats.html).

## Supplemental Item Titles

Supplemental Table 1. 20 informative genes for clusters (related to Figure 1)

Supplemental Table 2. CRISPR-screen results (related to Figure 2)

Supplemental Table 3. Stringent hit list DEV axis Stroma or TEC compared to iPSC+ (related to Figure 2)

Supplemental Table 4. Hit list DEV axis Stroma or TEC compared to iPSC+ (related to Figure 3)

Supplemental Table 5. Genes that score in TECs and/or Stroma (DEV axis) and have a kidney related human phenotype association (related to Figure 4)

Supplemental Table 6. CAKUT genes (human and mouse) that score in whole organoids (d35), TECs or Stroma (related to Figure 4)

Supplemental Table 7. Hits compared to established ciliary genes assembled by Reiter and Leroux, 2017 (related to Figure 5)

Supplemental Table 8. Potential novel ciliopathy genes: Ciliary genes leading to kidney phenotype in CRISPR screen (related to Figure 5)

Supplemental Table 9. KO leading to more tubular proliferation in both axis with metascape pathway enrichment analysis (related to Figure 6)

Note the different tabs in that file.

Supplemental Table 10. sgRNAs used in this paper (related to Figures 1, 5, 7)

